# Self-organising coordinate transformation with peaked and monotonic gain modulation in the primate dorsal visual pathway

**DOI:** 10.1101/327288

**Authors:** Daniel M. Navarro, Bedeho M. W. Mender, Hannah E. Smithson, Simon M. Stringer

## Abstract

We study a self-organising neural network model of how visual representations in the primate dorsal visual pathway are transformed from an eye-centred to head-centred frame of reference. The model has previously been shown to robustly develop head-centred output neurons with a standard trace learning rule [1], but only under limited conditions. Specifically it fails when incorporating visual input neurons with *monotonic* gain modulation by eye-position. Since eye-centred neurons with monotonic gain modulation are so common in the dorsal visual pathway, it is an important challenge to show how efferent synaptic connections from these neurons may self-organise to produce head-centred responses in a subpopulation of postsynaptic neurons. We show for the first time how a variety of modified, yet still biologically plausible, versions of the standard trace learning rule enable the model to perform a coordinate transformation from eye-centred to head-centred reference frames when the visual input neurons have monotonic gain modulation by eye-position.

## Author summary

Coordinate transformations are an essential aspect of behaviour. For instance, sensory information encoded in the coordinates of the retina needs to be transformed to relevant coordinates for planning and movement. Particularly, head-centred coordinates are essential for accurate motor behaviours and required to compute more complex coordinate transformations for sensorimotor integration [2]. Head-centred coordinates are obtained by combining information about the the retinal location of visual stimuli and the position of the eye. Previous work did not address the influence of different forms of gain modulation by eye position, albeit a variety of forms being widely reported for several cortical areas. Here we show how a biologically plausible model that successfully self-organised head-centred responses [1,3] fails when the visual input units have a commonly observed form of eye-position gain modulation, i.e. *monotonic modulation*. Our work makes an important contribution to understanding how head-centred responses may develop in the brain through an unsupervised process of visually-guided learning using a set of more sophisticated, and yet still biologically plausible, learning rules when visual precursor neurons have monotonic eye-position gain modulation. Furthermore, our findings may be applied to a range of other coordinate transformations with sensorimotor integration of monotonically encoded motor variables [4,5].

## Introduction

Within the primate dorsal visual pathway, neurons encode the locations of visual objects in different body-centred frames of reference. For example, neurons at an early stage of processing encode the locations of objects within an eye-centred reference frame, while neurons in later stages may encode the positions of objects with respect to the head or hand which is more relevant for guiding motor actions. A key question is how the visual system learns to perform such coordinate transformations between different body-centered reference frames. The neural mechanisms supporting this form of transformation, critical for visually guided motor function, have long been studied in primates. A neural phenomenon which is thought to play a key role in coordinate transformations in the dorsal visual pathway is *gain modulation*. This refers to the modulatory effect that a particular bodily state or posture, like eye or head position, has on the firing rate of some visual neurons responding in a given reference frame. Gain modulation of neuronal firing responses has been discovered in multiple stages of the primate dorsal visual pathway.

Parietal area 7a [6,7], LIP [8] and PO [9] have visual neurons modulated by the position of the eye in the orbit. This form of gain modulation is thought to be involved in supporting the development of head-centred visual representations, which were later identified in the same areas [10–12]. The parietal reach region (PRR) has hand-centred visual neurons with eye-position gain modulation [5], which are thought to be involved in guiding reaching to target locations. In certain regions such as parietal area 7a and LIP, the gain modulation is usually a *monotonic* function of the relevant bodily posture, such as eye or head position. For example, eye-position modulation often takes the form of a linear or saturating function of eye-position, which multiplicatively modulates the underlying Gaussian retinotopic receptive field of a visually responsive neuron [6]. While in other areas, such as V6A, there can be a higher proportion of retinotopic visual neurons with peaked eye-position gain fields [13].

Contemporary understanding of the potential role of such gain modulation in coordinate transformation has been informed by the highly influential coordinate transformation model of [14], reviewed below (section Previous Neural Network Models). The majority of models studying coordinate transformation since that time [4,5,15–17] have been heavily inspired by this early work. These models set up the synaptic weight matrix by some form of supervised, error correction, learning algorithm which cannot be implemented in the cortex [18]. Such algorithms provide no biologically plausible learning hypothesis, and also produce circuits which violate Dale’s law, the widely accepted neuroanatomical fact that a given presynaptic neuron cannot be both excitatory and inhibitory across its efferent synapses [19].

Our laboratory has developed a biologically plausible neural network model that self-organises its synaptic connectivity during visual experience such that the model learns to transform eye-centred visual representations to head-centred representations [1,3]. The model can achieve this using purely associative local synaptic learning rules with no supervisory training signal - i.e. unsupervised learning. The model is able to self-organise its synaptic connectivity by exploiting the natural eye and head movements of primates. However, a limitation of previous studies with this model has been their reliance on incorporating retinotopic visual input neurons with responses that are modulated by only *peaked* eye-position gain fields. This is not biologically realistic because many retinotopic visual neurons in the monkey brain are found to have *monotonic* eye-position gain fields. In the new simulations presented below, we first show that the incorporation of retinotopic visual input neurons with monotonic gain fields leads to a failure of the model to develop head-centred output neurons. We then show how model performance can be substantially improved by employing a range of more sophisticated, yet still biologically plausible, learning rules.

Next we present a review of relevant physiological and behavioural data along with a discussion of some previous computational models in order to provide the context for the new simulation results discussed in this paper.

## Physiology

### Eye-Position gain modulation of retinotopic visual neurons in parietal cortex

Computer simulations carried out within our laboratory have demonstrated that an important precursor to the emergence of head-centred visual neurons in the parietal cortex is likely to be the presence of retinotopic visual neurons with responses that are gain modulated by the position of the eye in the orbit [1,3]. A number of experimental studies have previously confirmed the existence of such retinotopic visual neurons with eye-position gain fields in the monkey brain.

The work by [6] was the earliest demonstration of the influence of the position of the eye in the orbit on the responses of retinotopic light sensitive neurons in area 7a of the monkey parietal cortex. The authors studied the responses of light sensitive neurons using both an attentive task and an inattentive task. In both tasks, the position of the head was kept fixed while the animal was able to move its eyes. During the attentive task the monkey had a peripheral stimulus flashed in a particular eye-centred location whilst maintaining fixation at some gaze angle. Conversely, during the inattentive task targets were flashed on random locations of the screen whilst the monkey freely oriented its gaze. The authors reported that the position of the eyes in the socket affected the responses of neurons in area 7a of the inferior parietal lobule during both attentive fixation and under the inattentive condition. Although it was also found that the responses of a much larger proportion of neurons were modulated by eye position during the attentive task than the inattentive task. The first task, the attentive task, revealed that 61% of neurons had their responses to visual stimuli significantly changed by the eye position. The neuronal responses could be more than three times stronger when the eye position shifted by 20° towards the preferred direction (optimal eye position). The second task, the inattentive task, revealed that a significantly smaller proportion of examined neurons (10%) had their responses to visual stimuli significantly changed by the eye position. The precise interaction between the visual signal encoding the retinotopic location of the target and the eye position signal was later characterized by [7]. These effects were also observed in the lateral intraparietal area (LIP) by [8]. This later work described such gain modulated responses as a multiplicative interaction between a Gaussian retinotopic receptive field and a *monotonic* (planar) eye position modulation component.

The presence of more *peaked* eye position gain modulation has been observed in the parietal occipital area (PO) by [9] and [13]. In the more recent of these two studies, [13] designed an experimental task to investigate the proportion of retinotopic visual neurons in area V6A of the primate brain that are modulated by eye position with either peaked (non-monotonic) or planar (monotonic) gain fields. During the experimental task the monkey had 9 equally spaced fixation locations organised as a 3 × 3 grid, with the visual stimulus always presented at the fixation point. These authors found that approximately 56% of recorded neurons had their responses modulated by the position of the eye. Furthermore, 27% of the neurons with responses that were modulated by eye position had planar gain fields, whilst the remaining 73% had peaked gain fields. The authors also explored the influence of each type of gain modulation on a traditional neural network model of sensorimotor transformation proposed by [20]. Their main motivation was to understand the implications of the functional form of gain fields, i.e. peaked (non-monotonic) *vs* planar (monotonic), for sensorimotor transformations in reaching tasks. In particular, they investigated how the functional form of the gain fields affected the performance of the model proposed by [20] to transform eye-centred visual representations into head-centred representations. The authors found that the incorporation of planar rather than peaked eye-position gain fields led to reduced model performance, with the population of output neurons providing a less accurate representation of the location of the visual stimulus with respect to the head.

### Head-centred neural responses

A number of experimental studies have found visual neurons in the monkey brain that encode the locations of visual stimuli with respect to the head, i.e. within a head-centred frame of reference. The model simulations that we present below in section Results seek to explain how such head-centred neurons may develop in the brain.

[11] was the first experimental study to provide evidence of head-centred visual representations in the preoptic area (PO) of the macaque brain. A head-centred representation of visual space is assumed to be important for visually guided reaching and perceptual stability. In fact, [21] had previously predicted the existence of head-centred visual neurons in PO due to the presence of retinotpic visual neurons with eye-position gain modulation in areas 7a [7], LIP [8] and V3A [21], which are a key precursor to the development of head-centred visual representations [1,3]. In the experimental study of [11], a head-restrained monkey performed a series of fixations to different locations on a screen whilst a visual target was presented in various other screen locations for each fixation. This allowed the authors to assess how individual neuron’s activity depended on either the eye-centred or head-centred location of the visual target. It was reported that the receptive fields of 11% of recorded neurons were not completely eye-centred, with six of these neurons presenting clear head-centred receptive fields. The authors concluded that previous claim by [8] that there was no explicit head-centred neuronal representation of visual space was incorrect. Furthermore, head-centred neurons were suggested to originate from pooling the output of preceding eye-position gain-modulated retinotopic visual neurons.

The work of [10] was motivated by the behavioural requirement for the brain to be able to integrate visual and auditory signals, such as the sight of moving lips and the corresponding speech sound, within a common body-centred frame of reference. The authors’ investigation of the reference frames used to encode visual and auditory responses in area LIP revealed the first head-centred visual representations in this area. This contradicted the suggestion made by [8] that head-centred responses would not exist in area LIP. The proportion of neurons sensitive to visual target locations was 72%, whilst 51.4% of neurons were sensitive to auditory target locations. Moreover, 5% to 43% of neurons were simultaneously responsive to both visual and auditory target locations, depending on how responsiveness was defined. For visual neurons, 33% had eye-centred responses and 18% had head-centred responses. Regarding auditory neurons, 10% had eye-centred responses while 23% had head-centred responses. Neither the remaining 49% of visual neurons nor the remaining 67% of auditory neurons could have their responses classified as eye-centred or head-centred. In summary, it was found that both auditory and visual neurons had responses compatible with either eye-centred or head-centred frames of reference, although most neurons had complex responses that could not be classified into either of these categories. This was the first time that head-centred neuronal responses were indentified in area LIP.

## Previous Neural Network Models

[14] developed an early influemtial model that learned to transform an input representation consisting of the position of the eye in the socket and the retinotopic position of the visual target to an output representation consisting of the position of the visual target with respect to the head. Specifically, two-dimensional representations of the eye-position **e** and of the retinal location of the visual target **r** were used as input for the neural network model. The target output used to guide the error-based update of the synaptic weights during training was the head-centred location **h** of the visual target represented by **r** + **e**. The model thus utilised a supervised, multi-layer backpropagation of error network architecture, in which the output layer of the network was provided with an explicit training signal representing the current head-centred location of the visual target. Such a network architecture is not biologically plausible for a couple of reasons. Firstly, it is not clear where such a training signal representing the current head-centred location of the visual target might originate from in the brain. Secondly, a multi-layer backropagation network architecture requires that the afferent weights to the hidden layer be updated using an error signal based on the efferent synaptic weights from these hidden neurons to the output layer scaled by the corresponding errors in the output layer. Such a network architecture in itself is not biologically plausible. Nevertheless, [14] found that training such a network to transform independent eye position and retinal target position signals to head-centred coordinates led to hidden units in the intermediate layer developing retinocentric receptive fields with planar eye-position gain fields, very similar to those found in the posterior parietal cortex by [6].

A number of later models that simulate the development of head-centred visual neurons have also relied on some form of error correction learning. For example, the model described by [15] relied on a form of supervised global error correction learning that utilised an error term which is unlikely to be present in the cortex. The model of sensorimotor transformation by [16] also used a supervised error correction learning rule to modify the synaptic connectivity within the network. So, although supervised error correction learning does not appear to offer a biologically plausible way of modelling the development of head-centred visual neurons in the brain, it has nevertheless remained a popular approach.

In contrast to the above error correction network models, [20] developed a model that utilised associative learning. These authors investigated how retinotopic visual representations could be transformed into head-centred representations that are relevant to reaching tasks. It was hypothesised that the observation of our own motor movements during a reaching task would be used to develop a map between the eye-centred visual representation of the target and the head-centred motor representation of the movement required to reach the target. In other words, the input layer of the neural network model represented the visual signals generated by observing the reaching movements, whilst the output layer represented the corresponding motor movements that were performed. As with [14], the input representation consisted of the position of the eye in the socket and the retinotopic position of the visual target, while the output representation consisted of the head-centred target position. Hebbian learning was then used during training to successfully associate each eye-centred visual input representation of the target to the corresponding head-centred output motor representation. In contrast to [14], the model developed by [20] did not use a backpropagation of error network architecture or any other form of error correction. Instead, they used a more biologically plausible associative learning rule to modify the synaptic connectivity within their model. However, the model was still trained in a supervised manner. That is, the network still made use of an explicit training signal in the output layer representing the head-centred location of the visual target in order to guide learning without an adequate explanation of where such a signal might originate from in the brain.

[22] proposed a hierarchical neural network model of coordinate transformation that used associative learning to set up the synaptic connectivity but did not require a biologically implausible supervisory training signal as used by [20]. The model developed by [22] had the following two main processing stages. The first processing stage used signals representing the retinotopic location of the visual target and signals representing the position of the eye to learn head-centred visual representations. The second processing stage used head position signals coupled with the head-centred visual representations developed by the first processing stage to learn body-centred visual representations. Training consisted of continously shifting a visual target presented to the network whilst randomly varying the position of the eyes and the head. Both processing stages used competitive learning with an associative learning rule to self-organise conjunctive representations of its respective inputs, and then used competitive learning with a form of temporally associative learning rule to bind representations that occurred close together in time. Learning in the second processing stage only started after the first processing stage had finished learning head-centred representations. This allowed each processing stage of the model to self-organise either head-centred or body-centred representations. Most importantly, in contrast to the models developed by [14] and [20], learning in [22] did not require the use of a biologically implausible supervisory training signal to guide learning of the coordinate transformations. However, the model of [22] did not investigate how the functional form of eye position gain modulation, i.e. monotonic *vs* peaked, may affect the development of head-centred visual representations. This is the focus of the current work presented below.

## Our approach: A Biologically Plausible Unsupervised Self-organising Neural Network Model of Coordinate Transformation

Our laboratory has previously published a biologically plausible neural network model of the visually-guided development of head-centred visual neurons, which relies on associative learning rules and does not require a supervisory training signal to guide learning [1, 3]. Instead, our model utlises an unsupervised form of competitive learning that exploits the natural statistics of how primates move their eyes and head as they shift their gaze around their visual environment. Specifically, the model employs a form of *trace learning*, which encourgages neurons in higher network layers to bind together input patterns that tend to occur close together in time. If a primate tends to move its eyes more frequently than adjusting the position of its head, then retinal images that occur close together in time will tend to correspond to different positions of the eyes but the same position of the head. In this case, trace learning will encourage neurons in higher layers to learn to respond to the position of a visual target in the same head-centred location across different retinal positions. Such neurons will have thus learned to represent the position of visual targets within a head-centred reference frame. The inputs to the model are eye-centred visual neurons that represent the locations of visual targets on the retina, but which have responses that are also gain modulated by the position of the eyes in the socket. In this paper, we investigate how the learning process depends on the functional form of this gain modulation by eye-position. Two forms of eye-position gain modulation are explored: monotonic gain modulation, which is dominant in most primate parietal areas (LIP, 7a, PRR), and peaked gain modulation which is primarily found in area **PO**.

In the simulations described below in section Results, the model is found to robustly develop head-centred output neurons with a standard trace learning rule when incorporating visual input neurons with *peaked* eye-position gain modulation [1], but not with *monotonic* eye-position gain modulation. Moreover, even if the model has its synaptic connectivity perfectly prewired to perform a coordinate transformation from eye-centred input neurons with monotonic gain modulation to head-centred output neurons, subsequently introducing plasticity into the synaptic connections with the standard trace learning rule quickly degrades the synaptic connectivity and eventually leads to elimination of the head-centred output responses. Since eye-centred visual neurons with monotonic eye-position gain modulation are so common in the dorsal visual pathway [7,23,24], it is an important challenge to show how efferent synaptic connections from these neurons may self-organise to produce head-centred visual responses in a subpopulation of postsynaptic receiving neurons. A subsequent analysis of the nature of the failure of the self-organisation of the synaptic connectivities led us to explore the performance of the model with a variety of modified, yet still biologically plausible, more powerful versions of the standard trace learning rule. The choice of the modified versions of the trace learning rule used in this paper was motivated by the superior performance of these learning rules reported by [25]. Here we show for the first time how these modified learning rules enable the model to learn to perform a coordinate transformation from eye-centred to head-centred reference frames even when the visual input neurons have monotonic gain modulation by eye-position.

## Materials and methods

### The Self-organisation of the Synaptic Connectivity within the Neural Network Model

The model uses four core components to self-organise head-centred visual representations through a biologically plausible process of visually guided learning. The first component is a population of input neurons that encode both the position of the eyes in the orbit and the retinotopic location of visual targets. Such retinotopic visual neurons with eye-position gain modulation have been identified in multiple primate cortical areas [7–9,23,24]. The second component is a population of output neurons that compete with each other through mutual inhibitory interactions, which is a standard feature of cortical architecture [18]. The third component is a local synaptic *trace learning* rule to update the feedforward synaptic connections between the input and output neurons. The trace learning rule is a local associative learning rule that incorporates an exponentially decaying temporal trace of past neuronal activity, and it has been widely used in the context of developing invariant visual object recognition [26,27]. The trace learning rule has the effect of encouraging individual postsynaptic neurons to learn to respond to subsets of input patterns that tend to occur close together in time. Finally, the fourth component comes under the assumption that visual stimuli are relatively static in a world reference frame during natural self-motion. This assumption is justified by experimental findings in which primates adjust their gaze more often by moving their eyes rather than the head itself [28]. This behavioural strategy to adjust gaze is preferable because it reduces the frequency of making more energetically costly and slow head movements. In fact, it has been found that during exploration of natural environments with free head, eye and body movements, at any time when there is movement, isolated head and isolated eye movements occurred 13.3% and 33.1% of the time respectively, whilst a mixture of movements was observed in the remaining time. That is, during natural movement there are periods when the eyes are moving whilst the head remains stationary with respect to the visual environment and visual objects also remain stationary within the environment.

These four model components allow the model to self-organise its synaptic connectivity during visually guided training in the following way. If the eyes move around a scene containing a stationary visual target while the head also remains stationary, then the visual system will receive a sequence of input patterns with the visual target in different retinal locations but the same head-centred location. That is, the visual target remains stationary in the head-centred space, but changes its position in the eye-centred space. The sequence of eye-positions and resulting retinal locations of the visual target are represented by retinotopic visual input neurons with responses that are gain modulated by eye-position. The synaptic trace learning rule is able to bind subsets of input patterns corresponding to a visual target in the same head-centred location, albeit with the visual target situated in different eye-centred locations, onto the same output neurons. This is because input patterns corresponding to a visual target situated in the same head-centred location tend to occur close together in time due to the statistics of natural eye and head movements, in which the eyes tend to saccade about a static visual scene while the head remains stationary. Moreover, the naturally rapid movements of the eyes may expose the visual system to many such input pattern sequences, where each such sequence has the visual target situated in the same head-centered location but different randomised subsets of retinal locations. This process will ensure that all possible input patterns corresponding to the same head-centred location but different retinal locations are eventually brought into temporal proximity with each other as training progresses. Hence, all input patterns corresponding to the same head-centred location but different retinal locations would tend to occur clustered together in time. This process continues with the visual target seen in a different position within the head-centred space every time the position of the head is shifted. That is, the natural head movements that occur between sequences of rapid eye movements would shift the location of the visual target to new head-centred locations. In this manner, the learning process is repeated with the visual target presented in many different head-centred locations. New subsets of output cells would learn to respond to the visual target in each different head-centred location due to the competitive interactions between the output cells. Consequently, the output layer would eventually develop neurons that cover the entire space of head-centred target locations.

### The Architecture of the Self-Organising Neural Network Model

The neural network architecture of the model is shown in Fig 1. The network consists of the following two layers of neurons. The first layer is a population of input neurons that simultaneously encode the eye-position of the agent and the retinal location of the visual target. These visual input neurons are modelled as retinotopic neurons with eye-position gain modulation. The eye and retinal position spaces, representing the range of eye-positions in orbit and retinal target locations, covered [−30°, 30°] and [−90°, 90°], respectively. Feedforward synaptic connections project from neurons in the input layer to neurons in the second layer.

**Fig 1.**
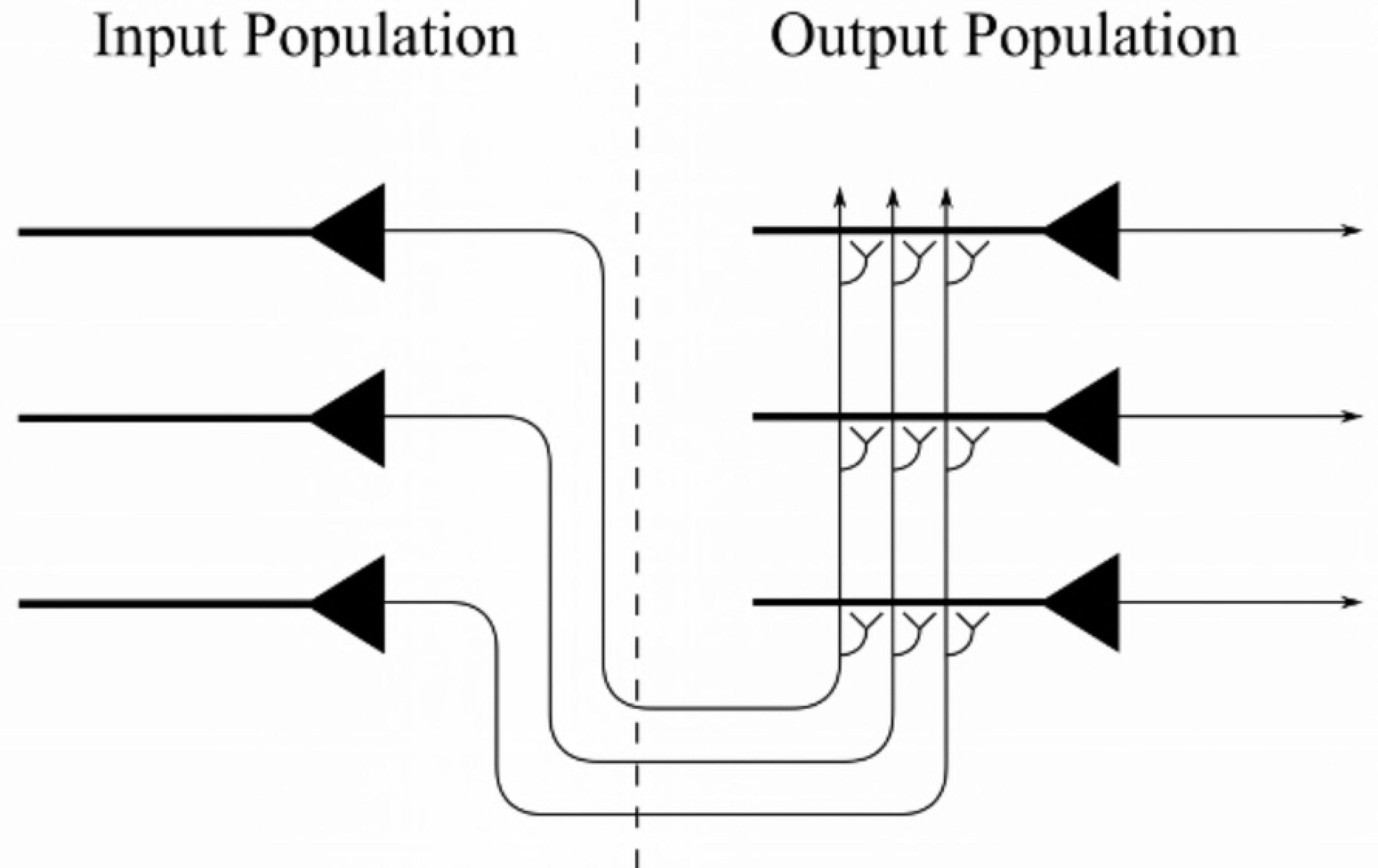
Architecture of the neural network model. The competitive output layer on the right receives afferent synaptic connections from neurons in the input layer on the left. A trace learning rule is used to modify the strengths of the feedforward synaptic connections from the input layer to the output layer during learning.

The second layer is a population of *N* output neurons that compete to represent patterns in the input layer [18]. Neurons in the second layer, the *output layer*, all receive the same number of afferent connections from neurons in the input layer, that is *ϕ* percent of the input population, but each output neuron receives connections from its own randomly assigned subset of the input neurons. Neither the output layer nor the input layer is topographically organised.

The strengths of the feedforward synaptic connections from the input layer to the output layer are initialised to random weights in the interval [0,1] at the start of each simulation. The synaptic weight vector of each output neuron is then renormalised as is typical in competitive networks [18].

### The Visually-Guided Training of the Network

The neural network is trained on input patterns that simultaneously encode both the position of the eyes in orbit and the retinotopic position of the visual target. The position of the eyes is kept within [−24°, 24°], whilst the retinal locations of visual targets are kept within the interval [−63°, 63°]. Keeping the position of the eyes and the retinal locations of visual targets within these respective intervals reduces edge effects due to clipping of the input representations.

Similarly, all M evenly spaced head-centred locations chosen for each experiment are confined within [−63°, 63°] to ensure that visual targets always remain in view as the eyes move. Each training epoch consists of *M* periods, where each period individually corresponds to one of the *M* chosen head-centred locations. During training, for each period a visual target is fixed in a given head-centred location whilst the eyes perform a series of saccades to *P* different and uniformly sampled eye-positions within [−24°, 24°]. The saccades between successive eye-positions are performed at a constant velocity of 400°/*s*. The duration of each fixation is set to 300*ms*.

Thus, the training process consisted of presenting to the network sequences of combined visual and eye-position input signals, which represent the visual target in fixed head-centred locations, whilst the eyes randomly shifted to different positions in the orbit.

### Testing the Network

The responses of the output units for all combinations of *T* head-centred visual target locations and *E* eye fixation positions are recorded after training to test the model. In order to test the ability of the trained model to generalise to new input patterns, the model is tested with a set of novel combinations of eye-position and visual target location not encountered during training. Specifically, the following *E* = 4 eye-positions are used during testing: −18°, −6°, 6° and 18°. For each of these eye-positions, the visual target is shifted in increments of 2° every 330ms to the next one of the *T* = 80 head-centred target locations within [−79°, 79°]. The firing rates of all output neurons are saved at the end of each fixation period to analyse the receptive field properties of the neurons, including the receptive field size, receptive field location and reference frame of response.

### The Neuronal and Synaptic Dynamics of the Model

#### Input Layer

The firing rates of neurons in the input layer were modelled by a response function that encodes the retinotopic location of a visual target, where the responses were modulated by the position of the eyes in the orbit. The response function of each input neuron *j* maps the current retinal location *r* of the visual target and the eye-position e onto the neuron’s instantaneous firing rate 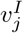. The instantaneous firing rate values are defined within the range [0, 1]. We investigated the performance of the model when the response functions of the input neurons were modulated by two different functional forms of eye positon gain field as described next.

The first response function has a peaked eye-position gain field as shown in Fig 2**A**. This form of eye-position modulation has been reported in cortical area PO by [9]. The full response function is described by

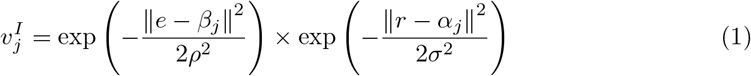

**Fig 2.**
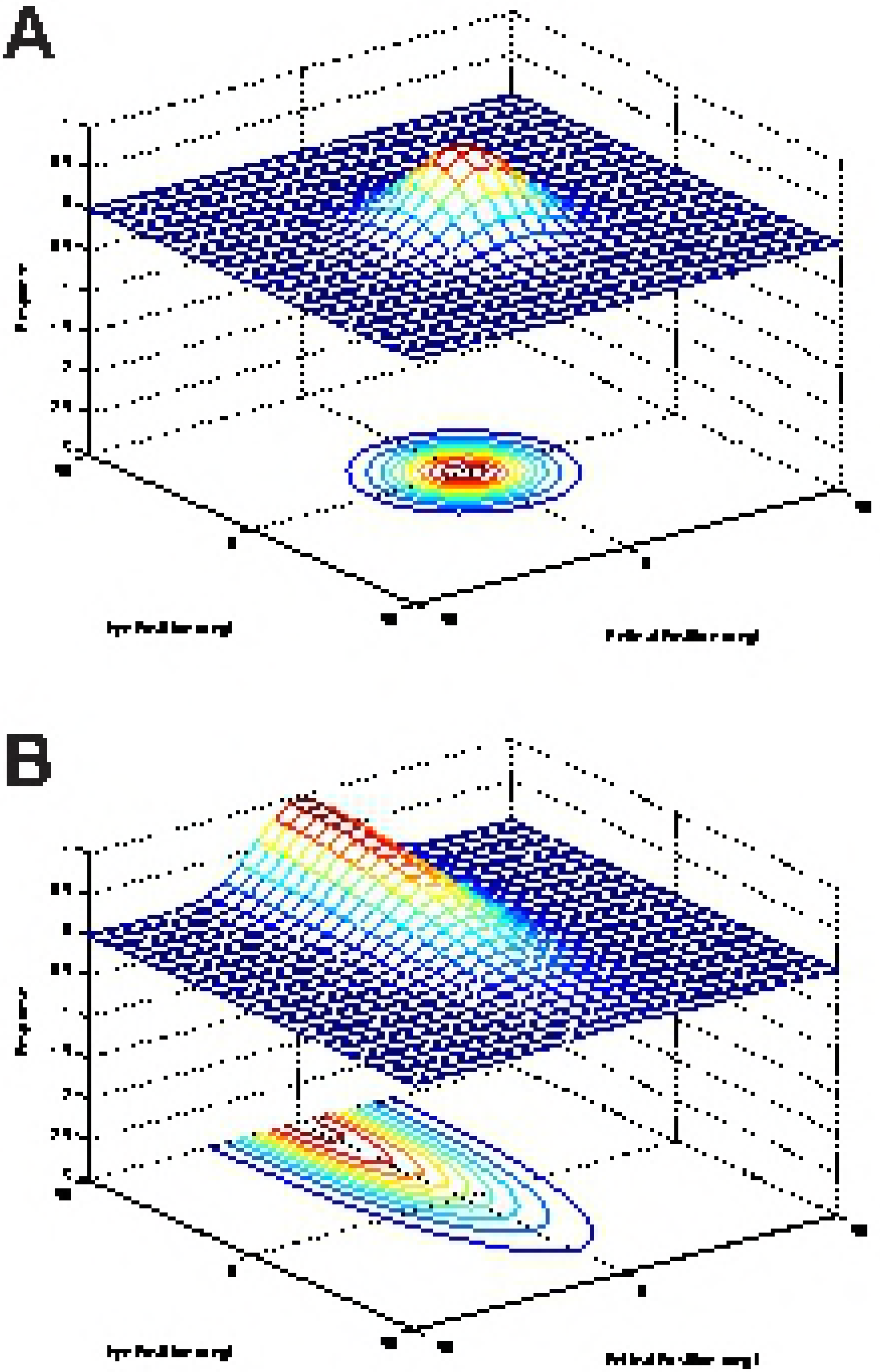
Examples of the two alternative response functions used to compute the firing rates of input neurons. For each type of response function, we plot the responses of an individual input neuron as a function of eye-position and retinal location of a visual target. A: Example of a peaked response function, in which the firing rate is modulated by a peaked eye-position gain field as described by Eq 1. B: Example of a monotonic response function that is modulated by a sigmoidal eye-position gain field described by Eq 2.

This response function is formed from a product of two components: the first component represents the eye-position signal, whilst the second component encodes the retinotopic position *r* of the visual target. In Eq 1, the neuronal response is modulatec by a peaked Gaussian function of eye-position. The parameter *β_j_* denotes the preferred eye-position for each input neuron *j*. The width of the corresponding Gaussian eye-position tuning curve is determined by the standard deviation *ρ*. The preferred retinal location of a target stimulus for each input neuron *j* is denoted by the paramete *α_j_*. The width of the corresponding Gaussian retinal tuning curve is determined by the standard deviation *σ*. Each input neuron *j* is set to respond maximally to a unique combination of retinal target location (*α_j_*) and eye-position (*β_j_*). The entire two dimensional space consisting of possible combinations of eye-position and retinal target location is covered by the population of input neurons in integer steps of 1 degree in each dimension. This results in a total of 201 × 61 = 12, 261 neurons in the input layer.

The second form of firing rate response function used to model the input neurons incorporates a sigmoid eye-position gain field as shown in Fig 2**B**. This form of eye-position modulation is monotonic in the eye-position dimension, as has been observed in multiple visual areas [7,8]. Although the form of the modulation is sigmoidal whilst most empirical work has described it as planar, [16] showed the data i also compatible with a saturating sigmoidal gain formulation. The full response function is described by

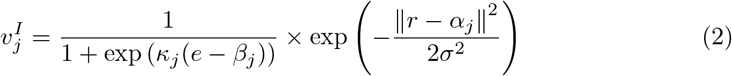

In Eq 2, the visual receptive field is modulated by a sigmoidal function of eye-position. For each input neuron *j*, the parameter *κ_j_* determines the gain direction and saturation rate of the modulation, where *κ_j_* is −2× the slope of the sigmoid. The inflection point *β_j_* determines the eye-position where a firing rate response greater than 0.5 begins. The input neurons all have the same absolute saturation rate, that is |*κ_j_*| = |*κ_m_*| for all *j* and *m*, but one half has a positive gain direction (*κ_j_* > 0) whilst the other half has a negative gain direction (*κ_j_* < 0). Each input neuron *j* is set to respond maximally to a unique combination of retinal target location *α_j_*, eye-position *β_j_*, and gain direction and saturation rate *κ_j_*. The population of input neurons evenly covers the entire three dimensional space resulting from such combinations. This results in an input population of 201 × 61 × 2 = 24, 522 neurons.

#### Output Layer

Three dynamical quantities were defined for each neuron *i* in the competitive output layer: an internal activation *h_i_*(*t*), a memory trace value *q_i_*(*t*), and an instantaneous firing rate *v_i_*(*t*) [29].

The internal activation is governed by the equation

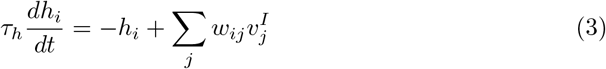

where *w_ij_* denotes the strength of the synapse from the *j*^th^ input neuron to the *i*^th^ output neuron and *τ_h_* is a time constant common for all output neurons.

The instantaneous firing rate is given by

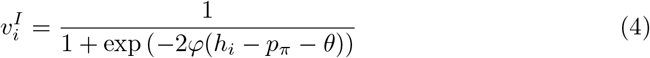

where *θ* and *φ* denote the sigmoid threshold and slope, respectively. The level of competition between neurons in the output layer, and thereby the proportion of neurons that remained active, is controlled by the parameter *p*_π_. Specifically, the parameter *p*_π_ is set to the π^th^ percentile point of the distribution of neuronal activations within the output population. For example, *p*_π_ is set to the top tenth percentile activation value when π is set to 90. This way of implementing competition within the competitive output layer has been previously used in competitive neural network models of the primate visual system with trace learning [30]. In the cortex, lateral inhibition is implemented via inhibitory interneurons [29]. The trace value *q_i_*(*t*) is defined in the following section.

#### Modification of Synaptic Weights by Trace Learning

Trace learning rules for synaptic modification encourage postsynaptic (output) neurons to bind together subsets of input patterns that tend to occur close together in time by incorporating a temporal trace of recent neuronal activity. The trace value *q_i_*(*t*) for the *i*^th^ neuron in the output layer is given by

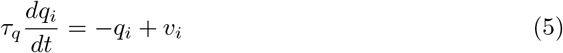

where *v_i_* is the instantaneous firing rate of the neuron, and *τ_q_* is a time constant common for all output neurons.

In the first part of the paper, during training the strength of the synapse from the *j*^th^ input neuron to the *i*^th^ output neuron is governed by the standard trace learning rule previously implemented by [1]

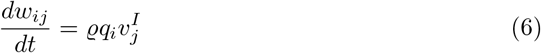

where *ϱ* is the learning rate, 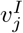 is the instantaneous firing rate of the *j*^th^ input neuron and *q_i_* is the trace value of the *i*^th^ output neuron. However, later in this paper we will introduce a number of new, more powerful forms of trace learning, which are in fact needed to produce head-centred output neurons when the input neurons are modulated by a sigmoidal (monotonic) function of eye-position.

Finally, after each weight update during training, the length of the weight vector for each output neuron *i*, that is 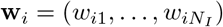 where there are *N_I_* input neurons, is renormalised by setting

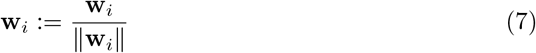

This prevents unbounded growth of the synaptic weights during training [29]. Experimental evidence for renormalisation of synaptic weights in the brain has been reported by [31].

### Simulation of the Differential Model

The Forward-Euler scheme is used to numerically integrate the coupled differential equations given by Eq 3 and Eq 5–6. The numerical time step Δ*t* is set to one tenth of the neuronal time constant *τ_h_*. We checked that the simulation results remained similar if the time step was reduced or the number of training epochs increased.

During training and testing, the input patterns encoding the changing eye-position and retinotopic target location are simulated dynamically and sampled at 1kHz. Linear interpolation is used to compute the numerical inputs to the discretised Forward Euler model equations, which require input values at every numerical time step Δ*t* = *τ_h_*/10.

### Analysis of Network Performance

In order to analyse whether individual output neurons are predominantly responding in an eye-centred or head-centred frame of reference, we used a method of analysis originally developed by [1]. This analysis is described next with the mathematical details taken from that earlier paper.

Let **R** be a matrix containing the responses of a given neuron during testing, where **R**[*i*, *j*] denotes the firing rate when the model is fixating in the *i*^th^ eye-position *e_i_* and the visual target is in the *j*^th^ head-centred location *t_j_*, as recorded during the testing protocol described above. The vector (**R**[*i*, 1],…, **R**[*i*, *T*]) is referred to as the response vector at the *i*^th^ eye-position. The number of eye-positions during testing is denoted by *E*, while the number of head-centred locations for visual targets during testing is denoted by *T*. The indexing of eye-positions and head-centred target locations are ordered from left (negative) to right (positive), that is *e*_1_ ≤ … ≤ *e_E_* and *t*_1_ ≤ … ≤ *t_T_*.

To determine which reference frame an output neuron is responding in during testing, two separate metrics are applied that reflected to what degree the neuronal response is compatible with either an eye-centred or head-centred reference frame, and then the values of these two metrics are compared.

The head-centredness metric computed the degree to which the head-centred response vectors of a neuron remained stable across different eye-positions. The head-centredness metric measured the degree of such stability for a given output neuron by averaging correlations between response vectors for different eye-positions, that is

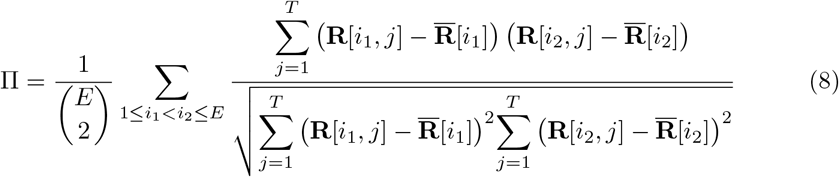

where

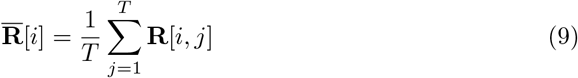

This yielded a metric which is referred to as the *head-centredness* of the output neuron, and it is bounded between −1 and 1, where a perfect correlation of 1 indicated a perfectly head-centred response.

A very similar analysis is done to quantify the compatibility of the responses of the output neuron with an eye-centred frame of reference. That is, a visual neuron is judged to respond in an eye-centred frame of reference to the extent that its eye-centred response vectors remain stable across different eye-positions. The eye-centred analysis proceeded as follows. To reiterate, each response vector (**R**[*i*, 1],…, **R**[*i*, *T*]) is the result of testing over the same set of head-centred locations, but with the model fixated in a distinct eye-position. Therefore, each response vector also corresponded to a unique range of retinal locations. The *intersection* of these retinal ranges corresponded to different portions of each response vector, and it is these portions that are subject to correlation analysis. Specifically, *f_i_* denotes the first vector position in the *i*^th^ response vector to be included, and the *V* − 1 next positions are included as well such that the subvector (**R**[*i*, *f_i_*],…, **R**[*i*, *f_i_* + (*V* − 1)]) is the vector being used for the correlation analysis. The derivation of *f*i** and *V* are found in the appendix Appendix A. This gave the metric

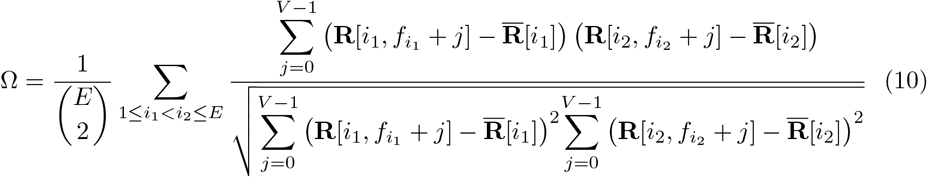

where

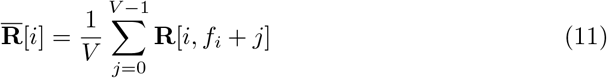

This is referred to as the *eye-centredness* of the output neuron, and it is bounded between −1 and 1, where a perfect correlation of 1 indicated a perfectly eye-centred response. Response vectors which had no response for the extracted ranges are excluded from the correlation, and a neuron without a response within this range of retinal locations at any eye-position is excluded from further analysis.

A neuron is finally classified as head-centred if П > 0 and П > Ω, and classified as eye-centred if Ω > 0 and Ω > П. If neither of these conditions is met then the neuron remains unclassified.

## Results

### Self-organisation with peaked and monotonic gain fields

This experiment explores the feasibility of the self-organisation of head-centred receptive fields under the two different forms of eye-position gain modulation. Two models are trained and tested on the same stimuli, where one model has peaked eye-position modulation in the input population as shown in Fig 2**A**, and the other model has sigmoidal modulation as shown in Fig 2**B**. The training lasts for 20 epochs. During each training epoch, a visual target is presented for approximately 5*s* in each of the eight head-centred training locations: −63°, −45°, −27°, −9°, 9°, 27°, 45° and 63°. For each period where the visual target is in a fixed head-centred target location, the eye-position is varied continuously through time as the model makes a series of saccades and fixations. During each such period, the model performs 14 saccades interleaved with 15 fixations, where each fixation lasts 300*ms*. Each saccade is at a constant velocity of 400°/*s*, and it is directed to a random eye-position within the range [−24°, 24°]. Each training epoch thus lasts for approximately 40*s*, and the entire training of the network is completed after about 800*s* of simulated time. The model is tested as previously described. The parameters for the two model simulations are given in Table 1.

Fig 3 compares the firing rate responses of the output neurons before and after training in the two models with either peaked or sigmoidal gain modulation of the visual input neurons. The responses of an output neuron from the model with peaked eye-position gain modulation before training (Fig 3**A**) exhibits no consistent structure in head-centred space across the different eye-positions. However, after training (Fig 3**B**) there is a clear maximal response to the same head-centred location across all eye-positions. Therefore, the self-organisation process has made the response reference frame of this output neuron strongly head-centred. The responses of an output neuron from the model with sigmoidal eye-position gain modulation before training (Fig 3**C**) also has an erratic and more eye-centred response prior to training due to the randomly assigned synaptic weights. However, unlike the peaked gain modulation model, training has the effect of making the neuron almost perfectly eye-centred. This is clearly seen by the receptive fields shifting in head-centred space in register with the eye-position shifts. Therefore, the self-organisation process has made this output neuron even more compatible with an eye-centred reference frame. The miniature scatter plots show the reference frame values of all neurons in the output layer, where each neuron is plotted as a point corresponding to that neuron’s particular combination of head-centredness and eye-centredness. The miniature scatter plots confirm the same general effects across the entire populations of output neurons. That is, subplot **(B)** shows that a large proportion of the output neurons in the model with peaked gain modulation have a high head-centredness and low eye-centredness, and thus respond in a head-centred reference frame. While subplot **(D)** shows that a large proportion of the output neurons in the model with sigmoidal gain modulation have a low head-centredness and high eye-centredness, and thus respond in an eye-centred reference frame.

**Table 1.** Simulation parameter of self-organising models with either peaked or sigmoidal eye-position gain modulation.

| Parameter | Symbol | Value (peaked) | Value (sigmoid) |
| --- | --- | --- | --- |
| Number of target locations | $M$ | 8 | 8 |
| Fixation sequence length | $P$ | 15 | 15 |
| Number of training epochs | - | 20 | 20 |
| Width of peaked eye-position tuning curve | $\rho$ | 6° | - |
| Width of retinal tuning curve | $\sigma$ | 6° | 6° |
| Gain magnitude | $\vert \kappa\vert $ | - | 0.0625 |
| Output neuron population size | $N$ | 900 | 900 |
| Input neuron population size | $N_I$ | 12261 | 24522 |
| Trace time constant | $\tau_q$ | 400ms | 400ms |
| Activation time constant | $\tau_h$ | 100ms | 100ms |
| Activation function slope | $\varphi$ | 4.5 | 4.5 |
| Activation function threshold | $\theta$ | 0.4 | 0 |
| Sparseness percentile | $\pi$ | 80% | 90% |
| Learning rate | $\varrho$ | 0.05 | 0.05 |
| Synaptic connectivity | $\phi$ | 5% | 5% |

**Fig 3.**
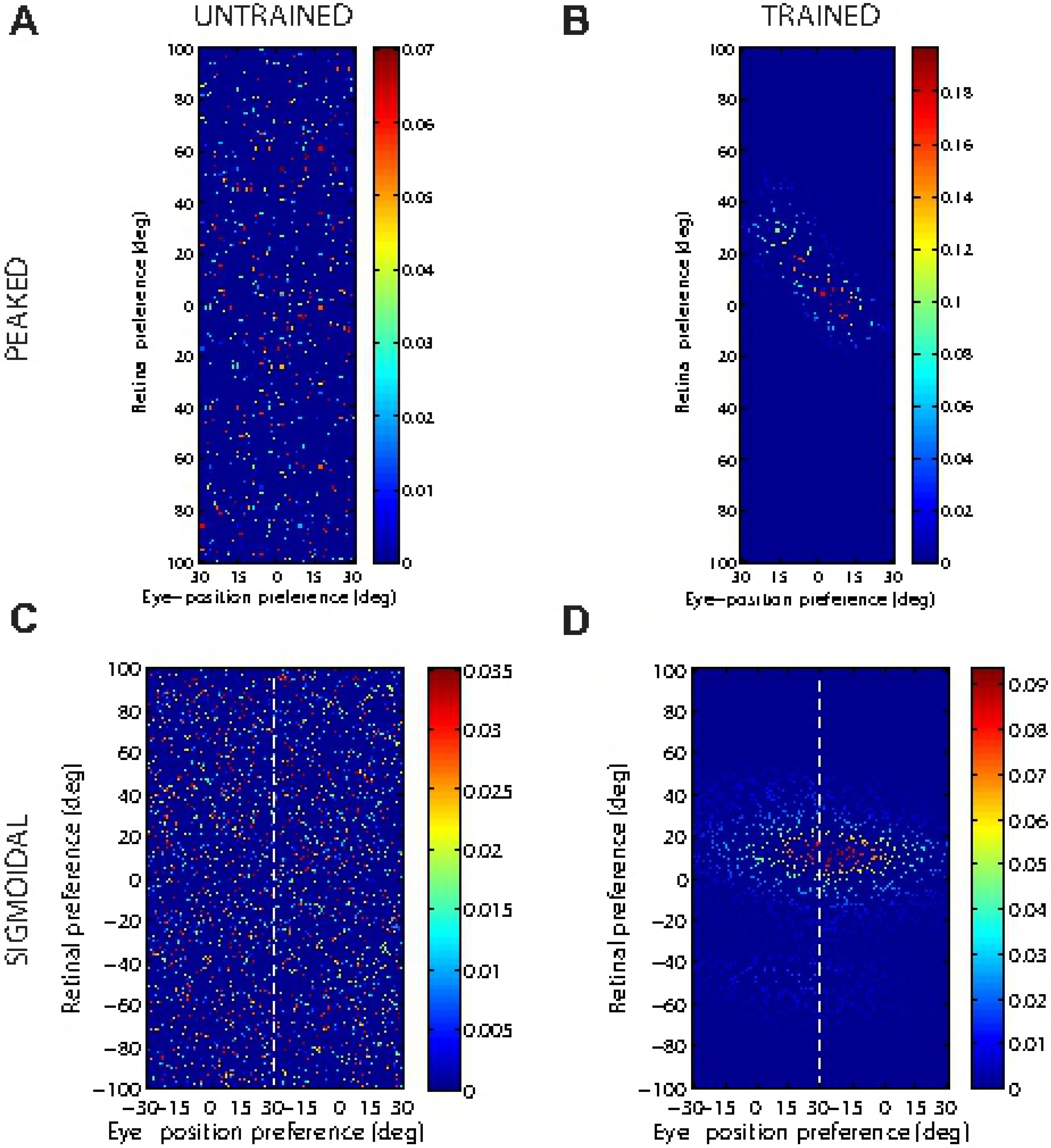
Comparison of model performance with peaked and sigmoidal (monotonic) gain modulation of visual input neurons. The figure shows the firing rate responses of output neurons before and after training with the standard trace rule (6). Specifically, each subplot shows the firing rate responses of a typical output neuron during testing for four different eye-positions: −18°, −6°, 6° and 18°. The top row shows output neuron #95 from the model with peaked eye-position gain modulation before training **(A)** and after training **(B)**. The bottom row shows output neuron #838 from the model with sigmoidal eye-position modulation before training **(C)** and after training **(D)**. In each subplot, each curve corresponds to a fixed eye-position while a visual target is presented across the same range of head-centred locations. It is evident in subplot **(B)** that after training the model with peaked gain modulation of the input neurons, the output neuron responds reasonably consistently when the visual target is presented within the localised interval of head-centred space [0°, 16°] regardless of the eye-position. The neuron is thus responding in a head-centred reference frame. However, in contrast, subplot **(D)** shows that after training the model with sigmoidal (monotonic) gain modulation, the responses of the output neuron in the head-centred visual space are much more dependent on the eye-position. Thus, this neuron is not representing the target position in a head-centred reference frame. The miniature scatter plots show the reference frame values of all neurons in the output layer, where each neuron is plotted as a point corresponding to that neuron’s particular combination of head-centredness (ordinate) and eye-centredness (abscissa). The neuron whose firing rate responses have been plotted is shown in the scatter plot by a red mark. The miniature scatter plots confirm the same general effects across the entire populations of output neurons. That is, subplot **(B)** shows that a large proportion of the output neurons are clustered in the top left quadrant of the scatter plot, indicating a high head-centredness (ordinate) and low eye-centredness (abscissa). These output neurons are thus responding in a head-centred frame of reference. While subplot **(D)** shows that a large proportion of the output neurons are clustered in the bottom right quadrant of the scatter plot, indicating a low head-centredness (ordinate) and high eye-centredness (abscissa). Thus, with monotonic gain fields acting on the input neurons, the population of output neurons have overwhelmingly learned to respond in an eye-centred reference frame.

Fig 4 shows the synaptic weight vectors of the same output neurons as those shown in Fig 3. Before training, there is no structure to the potentiated synapses of the output neurons in terms of the preferences of the presynaptic input neurons (Fig 4**A** and Fig 4**C**), reflecting the random weighting assigned to an untrained network. After training, the synaptic weight vector of the output neuron from the model with peaked eye-position gain modulation shows a clear diagonal structure (Fig 4**B**). This synaptic weight profile is consistent with a learned response to a particular location within the head-centred frame of reference, and is thus consistent with the observed head-centred responses of this neuron during testing (Fig 3**B**). The synaptic weight vector of the neuron from the model with sigmoidal (monotonic) gain modulation exhibites an entirely different pattern of potentiation after training. In this case, the strenghtened synapses have an approximately horizontal structure that is concentrated on input neurons corresponding to a small portion of retinal preference space (Fig 4**D**). As a result, this output neuron has learned an eye-centred response (Fig 3**D**).

**Fig 4.**
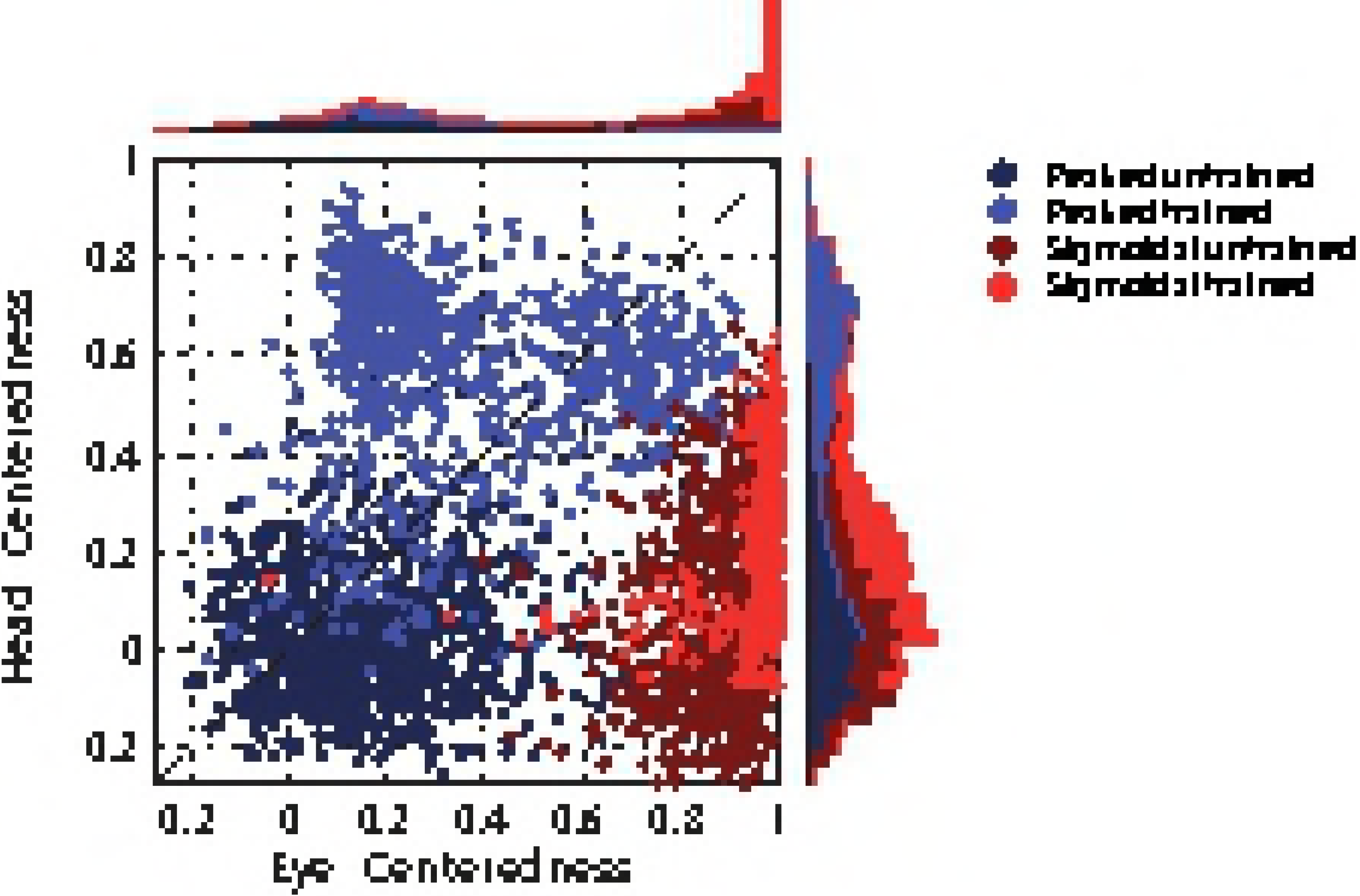
Strengths of the afferent synapses from the input population to a typical output neuron during testing. Results are shown for the model with peaked eye-position gain modulation before training **(A)** and after training **(B)**, and for the model with sigmoidal eye-position modulation before training **(C)** and after training **(D)**. The output neurons correspond to those plotted in Fig 3. In each plot, the afferent synapses have been arranged topographically by the preference of the input neuron for retinal location *α_i_* and eye-position **β_j_**. For the model with sigmoidal gain modulation, there are two input neurons for every combination of retinal preference and eye-position preference, but with opposite eye-position gain. Consequently, the input population has been separated by gain direction. The portion of each plot to the left of the white dashed line corresponds to input neurons with positive gain *κ_j_* > 0, while the portion of each plot to the right of the white dashed line corresponds to those input neurons with negative gain *κ_j_* < 0. It can be seen from subplot **(B)** that the output neuron in the trained network with peaked gain modulation has developed a diagonal weight structure, which is consistent with a learned response to a particular location within the head-centred frame of reference. In contrast, subplot **(D)** shows that the output neuron in the trained network with sigmoidal (monotonic) gain modulation has developed a more horizontal weight structure, which is consistent with a learned response to a specific location within the eye-centred reference frame.

Fig 5 shows the reference frame values for all output neurons from both models with either peaked or sigmoidal gain modulation tested before and after training. It is clear that, for the model with peaked eye-position modulation, training has the effect of making the majority of output neurons head-centred, and also with a much larger head-centredness value. Specifically, before training 26% of output neurons are head-centred, and after training 69% are head-centred. Moreover, among the head-centred neurons, the average head-centredness rose from 0.17 before training to 0.63 after training. For the model with sigmoidal eye-position modulation, training has the effect of keeping the majority of output neurons eye-centred, and indeed increasing their average eye-centredness from 0.88 to 0.96.

**Fig 5.**
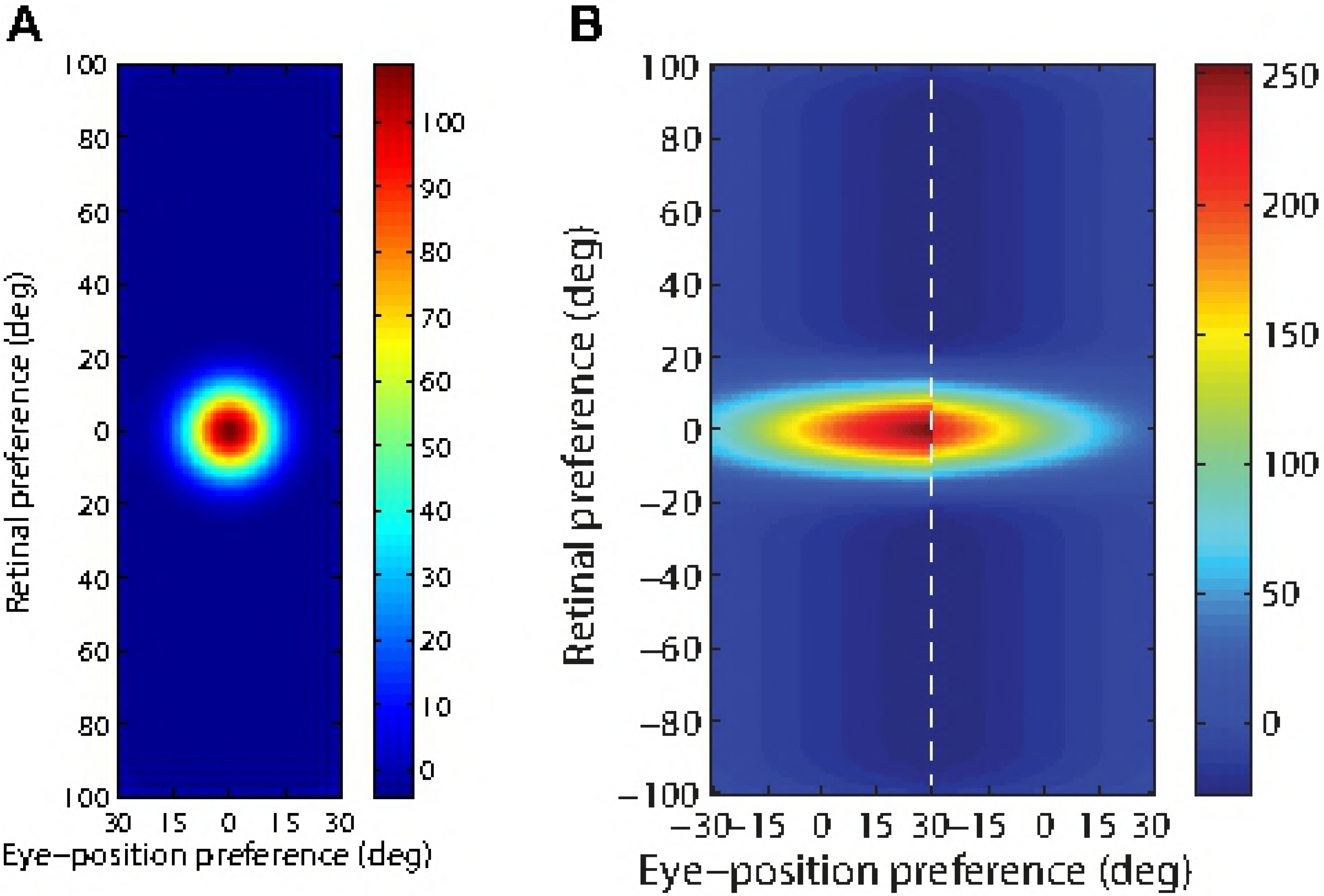
Scatter plot of eye-centredness and head-centredness values of output neurons from simulations with peaked and monotonic gain modulation. The scatter plot shows the eye-centredness and head-centredness values of all output neurons from four separate simulations corresponding to the models with peaked and sigmoidal (monotonic) gain modulation tested before and after training with the standard trace rule (Eq 6). Each point in the scatter plot corresponds to an output neuron from the given simulation, plotted in terms of its eye-centredness (abscissa) and head-centredness (ordinate). The dashed diagonal line with positive unity slope separates those neurons which are classified as head-centred (above line) from those that are classified as eye-centred (below the line). It is evident that after training most of the output neurons from the network with peaked gain modulation have become head-centred, while nearly all of the output neurons from the network with monotonic gain modulation have remained eye-centred.

In summary, when the visual input neurons have peaked eye positon gain modulation, training the network has the effect of developing head-centred output neurons. However, when the input neurons have sigmoidal (monotonic) gain modulation the training process makes most output neurons almost perfectly eye-centred.

Since a large proportion of visual neurons in the dorsal visual pathway have responses that are modulated by a monotonic function of eye-position, it is important to understand why monotonic gain fields make it more difficult for trace learning to produce head-centred output neurons. In the next section, we investigate this problem by carrying out a covariance analysis on the input patterns themselves.

### Covariance Analysis of the Effects of Gain Modulation

The preceeding model simulations failed to develop head-centred output representations during self-organisation when the input population had sigmoidal eye-position modulation, despite succeeding with peaked eye-position modulation. This raises the question of what the difference is between the two different forms of input encoding from the perspective of competitive learning [18]. It is well known that standard competitive networks develop weight vectors that reflect the covariance between the activities of input neurons. In particular, there is a tendency for output neurons to learn to respond to subsets of input neurons whose activities are highly correlated. Hence inspecting the covariance between input neurons across all input patterns may reveal what structure the weight vectors should converge towards under standard competitive learning conditions with a Hebbian learning rule.

The covariance between input neurons may be computed in the same way for both input neurons with peaked gain described by Eq 1 and input neurons with sigmoidal gain described by Eq 2. In the case of an input neuron with retinal preference *α* and eye-position preference *β*, the covariance between it and a second input neuron with corresponding preferences *α*\*, *β*\* is given by

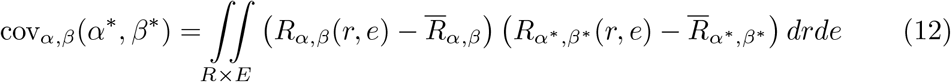

*R*_*α*,*β*_(*r*, *e*) is the response of a neuron with preferences *α*, *β* to a visual target at retinal location *r* and eye-position *e*, as given by either Eq 1 or Eq 2. The term 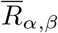 is the average response of the same neuron across all possible inputs in the *R* × *E* space, that is

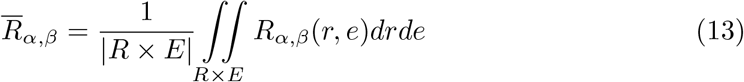

Fig 6**A** shows this covariance map for an input neuron with a peaked gain and preferences *α* = *β* = 0°, and Fig 6**B** shows the covariance map for an input neuron with sigmoidal gain and preferences *α* = *β* = 0° and *κ* > 0.

**Fig 6.**
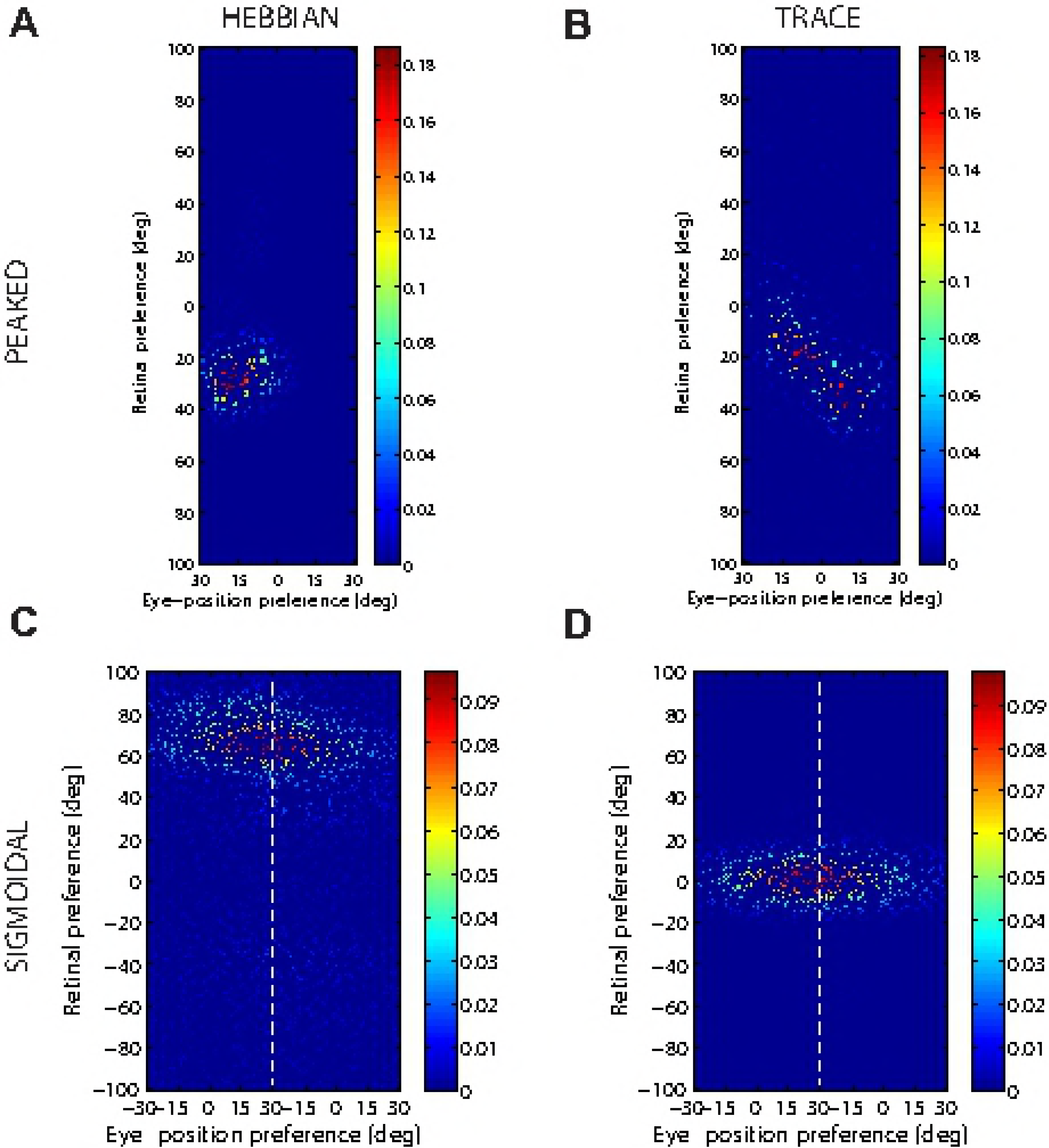
Covariance between a given input neuron and the rest of the input neuron population. Each plot shows the covariance between a given input neuron and the rest of the input neuron population in the form of a topographic map analogous to the weight vector maps shown above. Subplots **(A)** and **(B)** show results for input neurons with peaked and monotonic gain, respectively. In both cases the input neuron has preferences *α* = *β* = 0°, and in the monotonic case the input has positive gain (*κ* > 0).

The structure of covariance in the peaked modulation case is functionally identical to the response function of the input neuron, namely a two dimensional Gaussian tuning curve. The form of this covariance function is obvious by considering the correlations between the activities of input neurons with peaked gain.

In the sigmoidal (monotonic) modulation case, the situation is more complicated. The strong covariance is localised within the retinal preference dimension, but elongated within the eye-position dimension. This can again be understood by considering the response functions of the input neurons. Firstly, because all input neurons have a sharp, peaked tuning profile in the retinal preference dimension, any two input neurons need to have similar retinal preferences in order to have the possibility of being coactive. This explains the localisation of strong covariance in the retinal preference dimension. Secondly, the elongated form of the covariance function in the eye-position dimension results directly from the sigmoidal gain as follows. In the subpopulation of input neurons with a positive gain direction, similar to the reference input neuron (0°, 0°) itself, it is clear that other neurons with a similar retinal preference and with an eye-position preference to the right (i.e. larger than 0°) cofire more frequently with the reference neuron. This is because a positive gain implies that an input neuron responds to all eye-positions to the left of (i.e. smaller than) the eye-position preference of the neuron. Conversley, a negative gain implies that an input neuron responds to eye-positions to the right of (i.e. greater than) the eye-position preference of the neuron. Hence, in the subpopulation of input neurons with a negative gain direction, it can be seen that neurons with a similar retinal preference and with an eye-position preference to the left of (i.e. smaller than) 0° cofire more frequently with the reference neuron.

The covariance maps shown in Fig 6**A** and Fig 6**B** predict the structure of the weight vectors that we would expect to see develop in a competitive network with a standard Hebbian learning rule trained over all input patterns with either peaked or sigmoidal gain, respectively. These predictions are tested by running simulations with a Hebbian learning rule with both peaked and sigmoidal gain modulated input neurons. The Hebbian learning rule is implemented in the model by replacing the trace value *q_i_* in the standard trace learning rule (Eq 6) by the current firing rate *v_i_* of the postsynaptic neuron. For each simulation, there are 200 training patterns corresponding to random locations in the *E* × *R* space. The activation time constant is reduced to *τ_h_*= 30*ms* to avoid any trace effect during learning. Fig 7**A** and Fig 7**C** show the synaptic weight vectors of two typical output neurons that developed after training with the Hebbian learning rule when the input neurons are modulated by either peaked or monotonic gain, respectively. It is clear that these synaptic weight vectors have a very similar structure to the corresponding covariance maps shown in Fig 6. Thus, with the Hebbian learning rule, the underlying correlations between the activities of the input neurons with either peaked or sigmoidal gain shape the synaptic weight structure that develops during training. Most importantly, with monotonic gain, the synaptic weights are localised within the retinal preference dimension, but elongated within the eye-position preference dimension. This kind of synaptic weight structure leads to eye-centred output responses.

**Fig 7.**
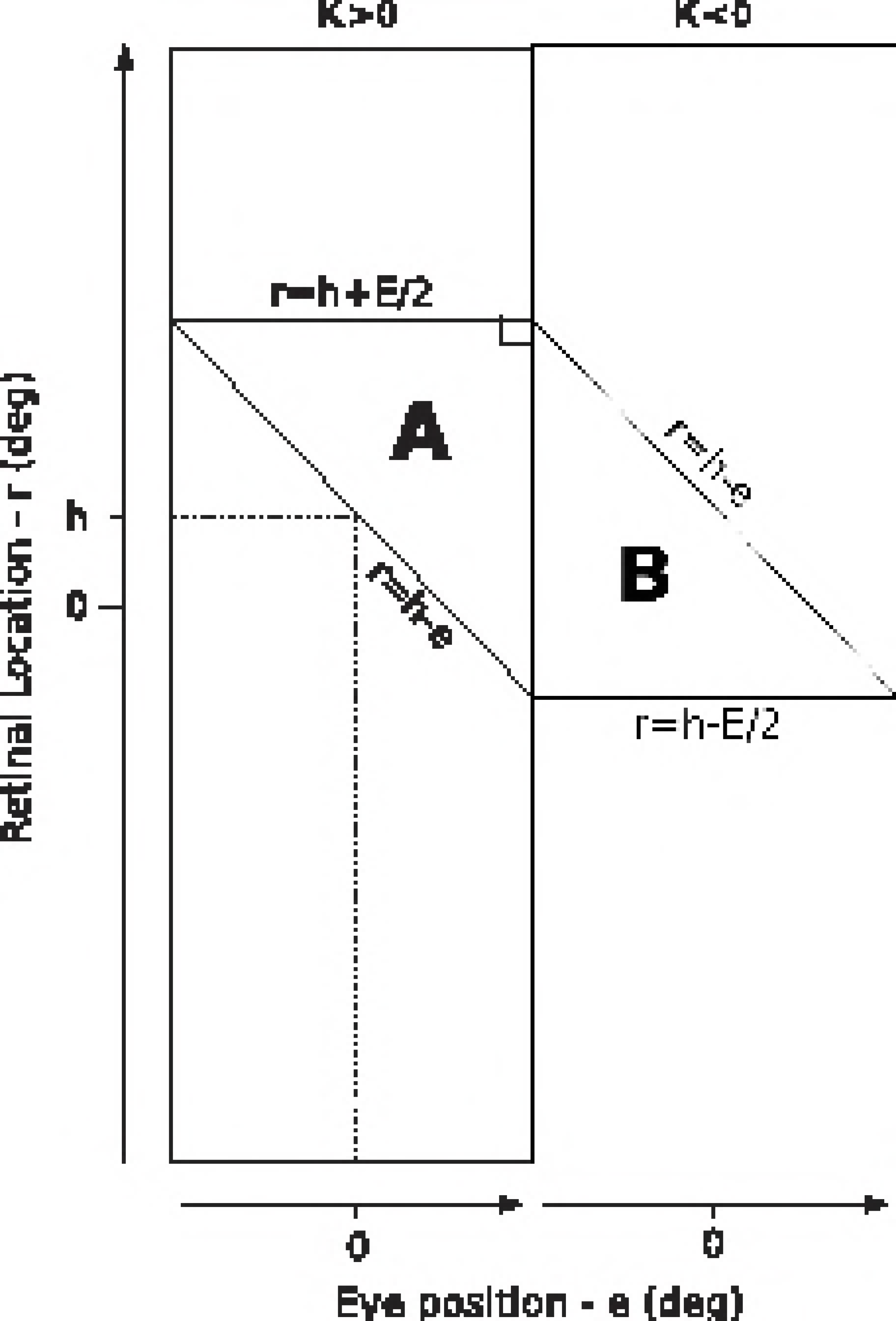
Weight vectors of two typical output neurons. The top row shows the weight vectors of two typical output neurons that develop when the input neurons have peaked eye-position gain modulation and the network is trained with either the Hebbian learning rule **(A)** or the trace learning rule **(B)**. The bottom row shows the weight vectors of two typical output neurons when the input neurons have monotonic eye-position gain and the network is trained with either the Hebbian learning rule **(C)** or the standard trace learning rule **(D)**.

Next, comparison simulations are run with the trace learning rule. Fig 7**B** and Fig 7**D** show the synaptic weight vectors of two typical output neurons that developed after training with the trace learning rule when the input neurons are modulated by either peaked or sigmoidal gain, respectively. Fig 7**B** shows a diagonal band of potentiated synaptic weights, which correspond to input neurons representing the same head-centred location but with different combinations of retinal location and eye-position. Thus, with peaked gain, the trace learning rule is able to simply bind together clusters of input neurons along a diagonal line in the (retinotopic preference × eye-position preference) input space corresponding to a particular head-centred location. Output neurons will then respond to particular head-centred locations regardless of eye-position or the retinal location of a visual target. However, the situation is quite different with sigmoidal gain modulated input neurons. Fig 7**D** shows a similar horizontal weight structure to that obtained with the Hebbian learning rule (Fig 7**C**). In particular, in both of these last two cases, the weight vector is very similar to the covariance structure found among the input neurons with sigmoidal eye-position modulation (Fig 6**B**). Thus, with sigmoidal gain, even if a trace learning rule is implemented, the output neurons still learn to represent eye-centred rather than head-centred locations. This is because developing head-centred output responses would require the trace learning rule to do more than simply bind input patterns together. With sigmoidal gain, trace learning must also disrupt and break apart output representations corresponding to clusters of highly correlated input neurons, which are localised in the retinotopic preference dimension but elongated in the eye-position preference dimension. However, in practice the standard trace learning rule given by Eq (6) is not strong enough to achieve this. Consequently, with the standard trace learning rule, these elongated clusters of input neurons with correlated activities continue to drive the development of eye-centred output neurons, as observed in the simulations previously reported in this article (section Self-organisation with peaked and monotonic gain fields).

### Introducing Plasticity into a Prewired Model

The failure of self-organisation to produce head-centred output neurons in a model with sigmoidal (monotonic) eye-position gain modulation suggests the following two important questions. First, does there actually exist a synaptic weight connectivity matrix that would support a mapping to head-centred output representations, even if in practice the self-organisation process using the standard trace learning rule (6) originally implemented by [1] fails to converge on this solution? Secondly, if it is possible to prewire a network with such a synaptic weight structure, then would the head-centred output representations be abolished by subsequently introducing synaptic plasticity either in the form of the standard trace learning rule (6) or normal Hebbian learning (where the trace value *q_i_* is replaced by the current firing rate *v_i_*)? If so, then this would demonstrate an even deeper problem: the presence of such forms of synaptic plasticity in a network with sigmoidal gain modulation not only fails to drive the development of head-centred output representations, but would also abolish any existing head-centred representations.

To address the above two questions, we construct a manually prewired model that is designed to produce head-centred output neurons with input neurons that are modulated by sigmoidal functions of eye-position. The prewired model is constructed as follows. There are 24522 neurons in the input population, each corresponding to a unique combination of retinal-position preference (*α_i_*), eye-position preference (*β_j_*) and slope (*κ_j_*). There are 900 neurons in the output population, each given a head-centred receptive field at one among nine head-centred locations, which are −68°, −51°, −34°, −17°, 0°, 17°, 34°, 51° and 68°. Each neuron in the output population is postsynaptically connected to a randomly assigned subpopulation of the input population. There are only two synaptic weight values across all synapses, simply referred to as elevated and depressed. The strength of a synapse is elevated if the presynaptic input neuron responded maximally to a combination of eye-position and retinal location corresponding to a head-centred location that is closest to the head-centred location assigned to the output neuron. Otherwise the synapse is depressed. Therefore, the output neuron receives strong driving input to the extent that a visual target is near its assigned head-centred location. Specifically, the weight of a synapse with a postsynaptic neuron assigned to head-centred receptive field location *h* and a presynaptic neuron having retinal-preference *α*, eye-position preference *β* and *κ* > 0 is given by

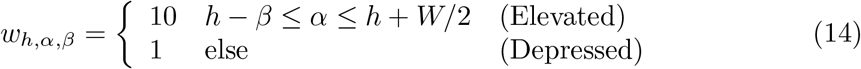

where *W* = 60° is the size of the eye-position dimension. Likewise when *κ* < 0 the weight is given by

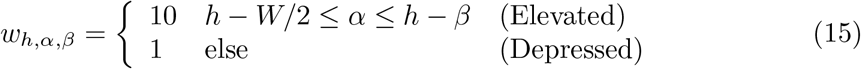

Fig 8 shows the structure of the canonical weight vector produced by Eq 14–15. Before testing the network, the synaptic weight vectors of all output neurons underwent the normalization step described by Eq 7. To provide a baseline for comparison, a network with randomly wired synaptic connections is also tested in the same way. The parameters for both models are given in Table 2.

**Fig 8.**
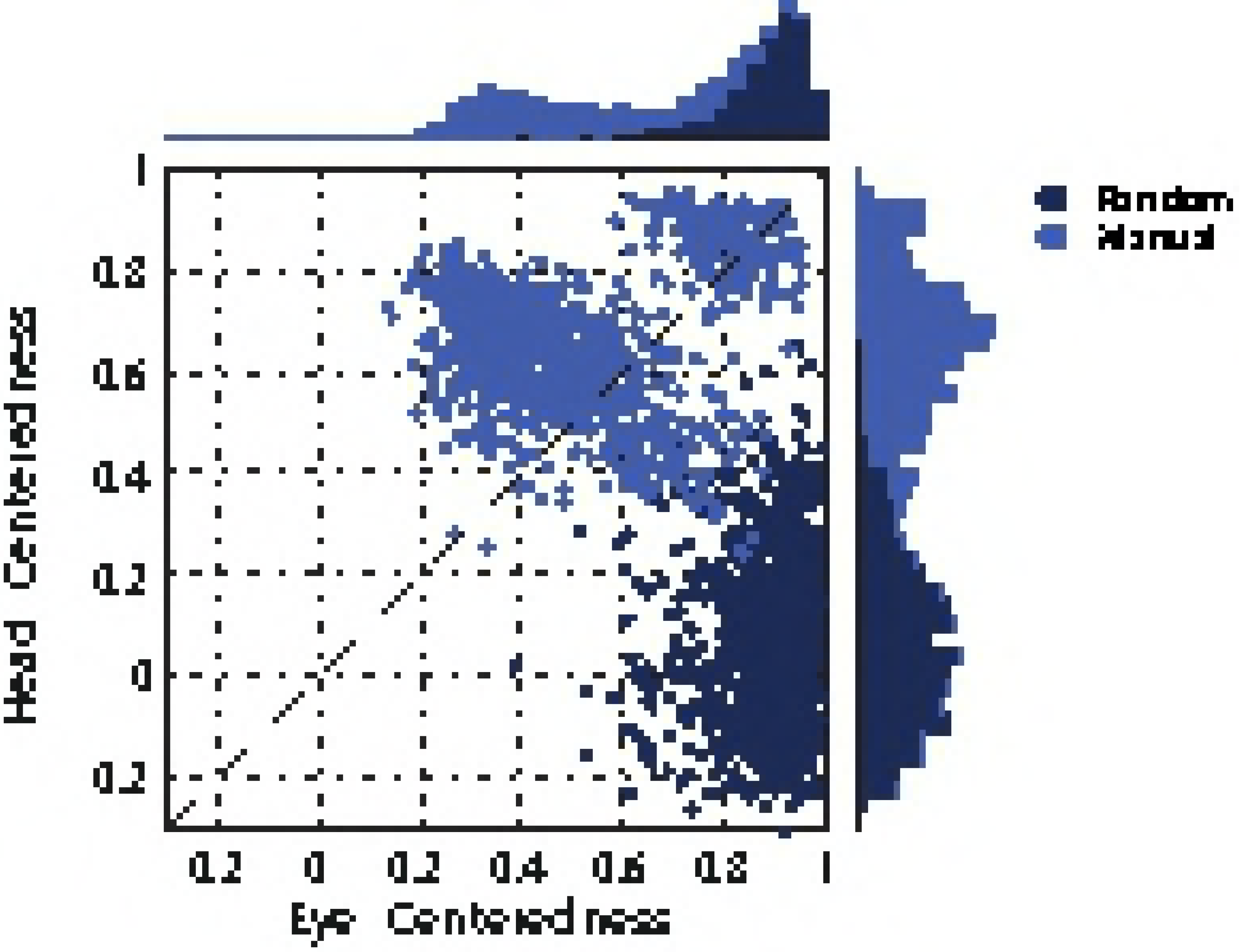
Synaptic weight structure of a network model that has been manually prewired in order to produce head-centred output neurons with input neurons that are modulated by a sigmoidal function of eye-position. The figure shows the structure of the canonical weight vector resulting from the prewiring Eq 14 and Eq 15. Each of the two rectangles represents the topographic organisation of one half of the input population in terms of retinal-preference (*α_i_*) and eye-position preference (*β_j_*), with the input neurons in the left rectangle having *κ* > 0 (positive gain) and the right rectangle having *κ* < 0 (negative gain). A neuron in the competitive output population which has been assigned a head-centred receptive field at location *h* will have elevated connections from input neurons with preferences located in the right-angled triangles of the input space, labeled **A** and **B**.

Fig 9 shows the eye-centredness and head-centredness values of output neurons in the manually prewired model as well as a randomly wired model for comparison. It is clear that in the manually prewired model, the majority of output neurons are head-centred. In contrast, there are no head-centred neurons in the randomly wired model. In summary, the results from the manually prewired model demonstrate the existence of a synaptic weight matrix which allows the output neurons to perform the desired coordinate transformation to a head-centred reference frame even when the input neurons are modulated by a sigmoidal function of eye-position.

**Table 1.**
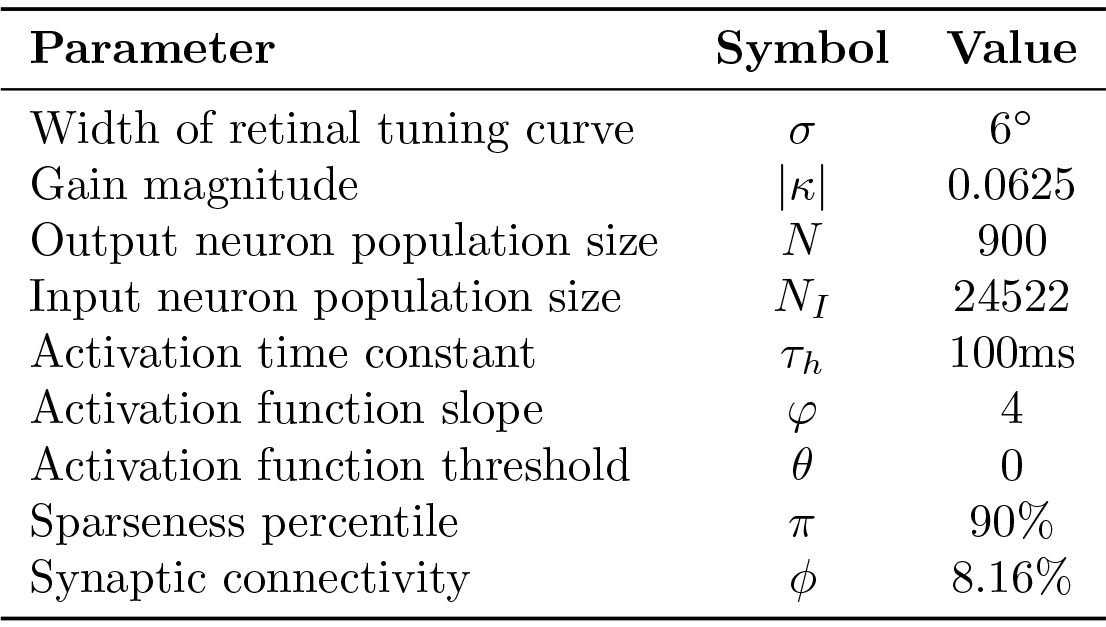
Simulation parameters of the network model that has been manually prewired in order to produce head-centred output neurons with monotonic modulated input neurons.

**Fig 9.**
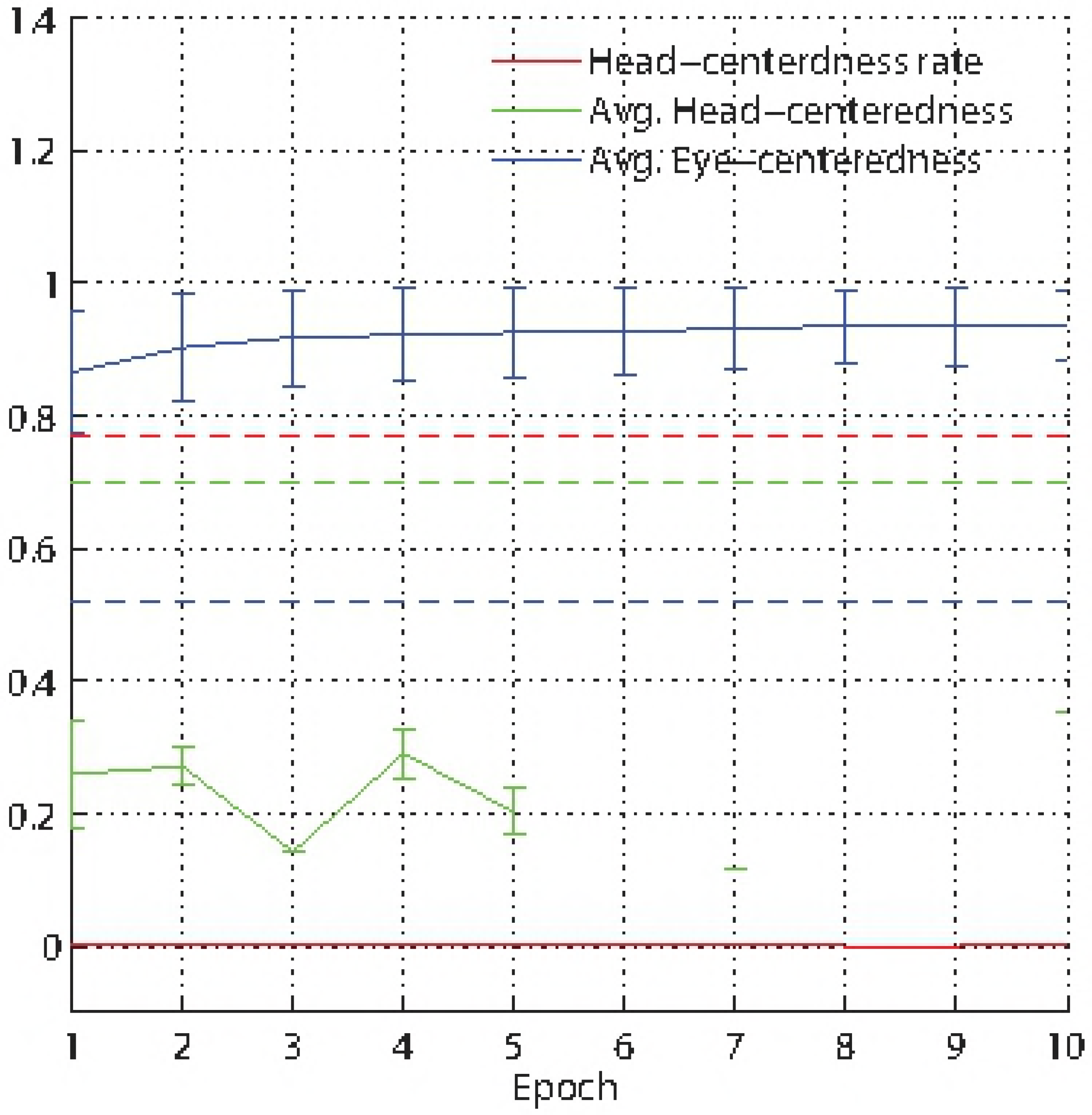
Performance of the prewired network model with monotonic modulated input neurons. The Figure shows the performance of the network model that has been manually prewired to produce head-centred output neurons with input neurons that are modulated by a sigmoidal function of eye-position. The scatter plot shows the eye-centredness and head-centredness values of all output neurons from the manually prewired model and a randomly wired model. Same conventions as in Fig 5. It can be seen that the majority of the output neurons in the manually prewired model display head-centred responses.

The next question to be explored is, what would be the effect of introducing synaptic plasticity, either in the form of the standard trace learning rule (6) or normal Hebbian learning, into the manually prewired model while it is exposed to the kinds of visual input described above? This experiment would inform whether these forms of plasticity not only failed to drive the development of head-centred output representations during self-organisation, but would even abolish existing head-centred representations. The manually prewired model is therefore subjected to the same visual training stimuli as described above over 10 training epochs. The results presented here are from a simulation using the standard trace learning rule (6). However, although not
shown, further simulations with a Hebbian learning rule without an explicit memory trace gave qualitatively similar results.

The impact of introducing synaptic plasticity into the manually prewired modelsas is inspected by plotting key summary statistics as a function of the number of training epochs in Fig 10. There is a catastrophic drop in model performance after only the first epoch of training, where the fraction of head-centred neurons decreased from ~77% to ~0.3%, and the average head-centredness among head-centred neurons decreased from ~0.7 to ~0.26. Subsequent training epochs remained around these levels. After epoch 5 there is not a single head-centred neuron.

**Fig 10.**
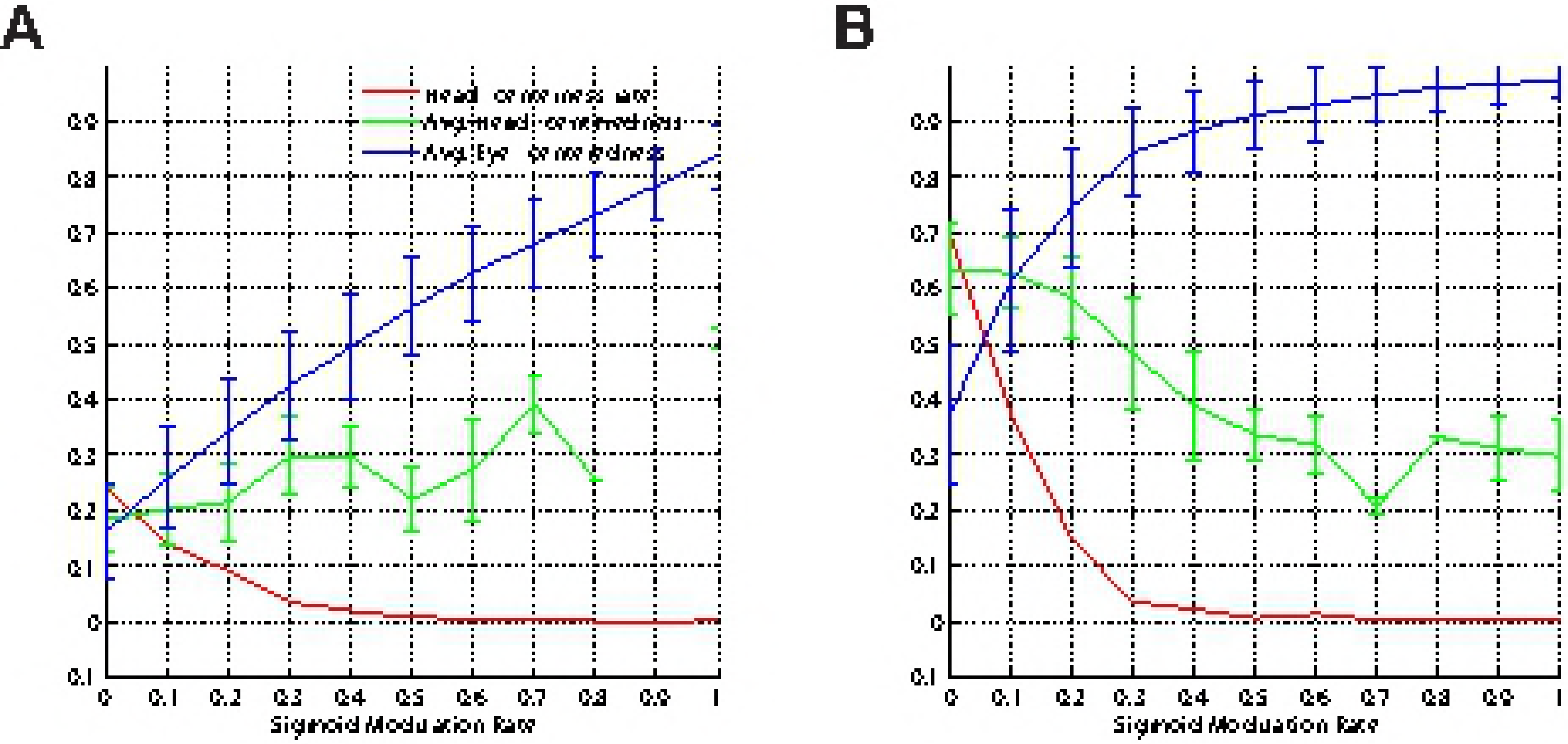
The effects of introducing synaptic plasticity into the network that has been manually prewired to produce head-centred output neurons when the input neurons that are modulated by a sigmoidal (monotonic) function of eye-position. The figure shows population analyses of the response properties of output neurons in the manually prewired model as the synaptic weights are further modified during ten training epochs with the standard trace learning rule (6). Three key summary statistics are given. The head-centredness rate (red) is the fraction of head-centred neurons in the output population. The average head-centredness (green) is the average head-centredness among head-centred neurons, and becomes undefined if no head-centred neurons are found to exist. The average eye-centredness (blue) is the average eye-centredness among all output neurons. The dashed lines show these values for the manually prewired network before training, while the unbroken lines show the values through successive training epochs after synaptic plasticity has been introduced. The error bars are the standard deviations. It can be seen that by the end of the first training epoch the majority of the output neurons switched from being head-centred to eye-centred.

In summary, it is found that just a single training epoch switched most of the output neurons from being head-centred to eye-centred. We hypothesised that this is due to the same visually-guided learning dynamics described above in section Self-organisation with peaked and monotonic gain fields and in section Covariance Analysis of the Effects of Gain Modulation, which come into operation when the retinotopic input neurons have monotonic eye-position gain modulation. Thus, even if the synaptic weights are initially manually prewired to effect head-centred output responses, which might be suggested to happen in the brain through genetic specification, the introduction of just a limited amount of synaptic plasticity, either in the form of the standard trace learning rule (6) or normal Hebbian learning, and visually-guided learning led to the output neurons rapidly switching to eye-centred responses. The presence of even modest levels of such synaptic plasticity will quickly overwrite head-centred representations that have been set up through structured (e.g. genetic) prewiring. Thus, since plasticity is ubiquitous in primate cortex, this suggests that any explanation for the development of head-centred visual responses must utilise a more sophisticated visually-guided learning process than demonstrated by [1], who considered only models with input neurons that were modulated by peaked functions of eye-position. Moreover, the loss of head-centred representations in the manually prewired model by introducing synaptic plasticity also represents a major challenge to the plausibility of previously published models, such as that of [20], which rely on an initial period of supervised learning to establish the required synaptic connectivity. The problem here is that when the supervisory training signal is eventually removed, the continued presence of associative plasticity may degrade and eventually abolish the head-centred output representations.

In the remainder of the paper we explore a variety of biologically plausible model variations that are aimed at discovering potential mechanisms by which head-centred output neurons may still develop through visually guided learning even when the network contains input neurons with monotonic modulation by eye-position. We begin by exploring the performance of the model when it incorporates a mixture of input neurons that are modulated by peaked and sigmoidal eye-position gain fields. After this, we explore the operation of the model with a number of more sophisticated, modified synaptic learning rules originally developed by [25] in the context of transform invariant visual object recognition, which maintain biological plausibility by continuing to rely on the locally available activities of the pre- and post-synaptic neurons. The choice of the modified versions of the trace learning rule used in the following sections of this paper is motivated by the superior performance of these learning rules reported by [25]. Finally, we conclude with the investigation of the performance of the model incorporating a mixed population of peaked and monotonic modulated visual input neurons with the synaptic weights also adjusted using a new modified version trace learning rule.

### Standard Trace Learning Rule with Mixed Peaked and Sigmoidal eye-position Modulation of Input Neurons

In this experiment it is investigated how mixing peaked and sigmoidal gain modulation in the input population in varying proportions would influence the development of head-centred output neurons in the self-organising model with the standard trace learning rule (6). This is an important issue since all cortical areas with eye-position gain modulation exhibit a mixture of different forms of modulation [7–9]. A series of simulations are conducted where each neuron in the input population is independently and randomly set to have either a peaked or sigmoidal gain modulation. Specifically, each input neuron is changed from having peaked to sigmoidal modulation with a probability *p*, called the sigmoid modulation rate, and values of *p* = 0,0.1,…, 1.0 are explored.

The impact of varying the sigmoid modulation rate on the characteristics of the model is inspected by plotting key summary statistics as a function of *p* in Fig 11. As expected, the head-centredness rate decreased as the sigmoid modulation rate increased, both in the trained and untrained models. However, as long as the sigmoid modulation rate is less than 30%, the trained model had a higher proportion of head-centred output neurons than the untrained model. In particular, for sigmoid modulation rates less than 20%, the fraction of head-centred neurons in the trained model did not drop below ~15%, and the average head-centredness among head-centred neurons remained no less than ~0.58.

**Fig 11.**
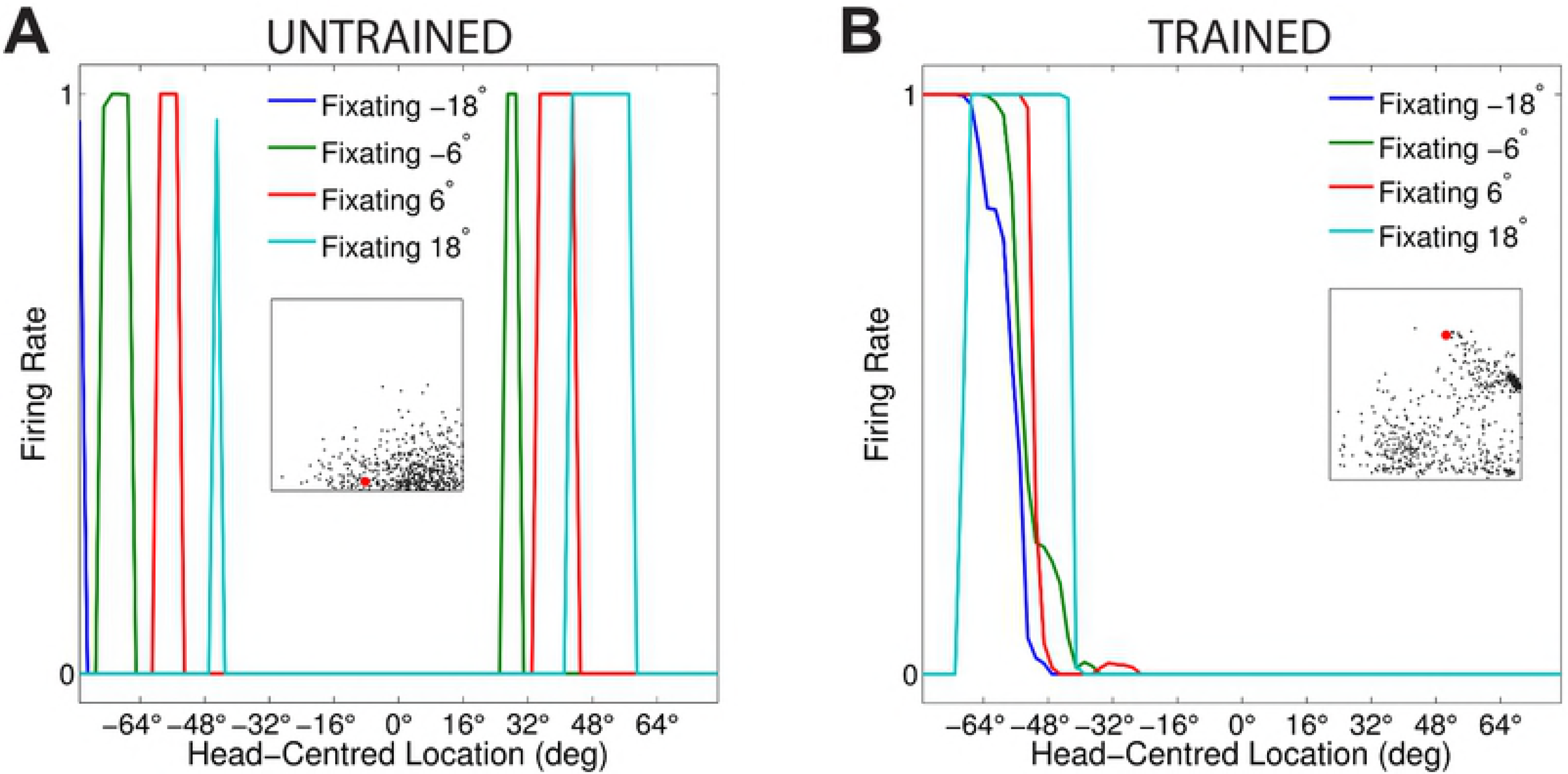
The effects of incorporating a mixed population of input neurons with both peaked and monotonic eye-position gain modulation. The plots show how the performance metrics vary with the monotonic modulation rate, *p*, which is the probability of each input neuron having a monotonic eye-position gain modulation. Results are presented showing the response characteristics of the output neurons before training **(A)** and after training **(B)**. Conventions are similar to Fig 10. It is evident that the head-centredness rate decreased as the sigmoid modulation rate increased, both in the trained and untrained models. However, as long as the sigmoid modulation rate is less than 30%, the trained model had a higher proportion of head-centred output neurons than the untrained model.

In summary, these results showed that when there is a large proportion of input neurons with peaked eye-position gain modulation, say with 0 ≤ *p* ≤ 0.2, then the self-organising model is still capable of developing a significant proportion, i.e. no less than ~15%, of head-centred ouput neurons during training. However, as the sigmoid modulation rate increased, the performance of the model deteriorated with far fewer head-centred output neurons present in the trained model.

Given that the model incorporating a mixed population of input neurons failed to develop a significant proportion of head-centred output cells whenever *p* > 0.2, we next investigated whether the introduction of more powerful, modified synaptic learning rules could produce head-centred output neurons when the entire population of input neurons were again modulated by a sigmoidal function of eye-position. Our choice of the modified versions of the trace learning rule adopted in this paper was motivated by their superior performance reported in previous work [25]. These modified versions of the standard trace learning rule are still biologically plausible in terms of only using locally available signals to update the synaptic weights of connections. Moreover, we expected that the introduction of a delayed trace of synaptic activity and as well as an anti-Hebbian component incorporated in these modified versions of the trace learning rule would provide the model with a way of weakening the observed Hebbian-like training behaviour evidenced, for example, by the comparison of Fig 7**D** and Fig 7**C**, and therefore to facilitate the self-organisation of head-centred responses with visual input neurons with monotonic eye-position gain modulation. The absence of these components makes other classic Hebbian-based learning rules (e.g. Oja’s rule [32]) ineffective in this case.

### Modified Learning Rule: Delayed Postsynaptic Trace with Anti-Hebbian Learning

[25] investigated how a set of modified more powerful versions of the trace learning rule can produce improved temporal binding and invariance learning. In particular, the authors showed that the performance of the trace learning rule is substantially improved by incorporating a trace of previous neuronal activity with an explicit time delay. This had the effect of removing the purely Hebbian term of the learning rule [25]. In the next simulations, the learning rules proposed by [25] were adapted to differential formulations for the time-continuous scenario in which the simulations are performed. A Forward Euler scheme was used to numerically integrate the differential equations. In all simulations the numerical time step was kept as one tenth of the neuronal time constant *τ_h_*. Unless explicitly mentioned, the learning rule was the only change from previous simulations.

This section presents simulation results showing the performance of a modified learning rule incorporating a delayed postsynaptic trace with anti-Hebbian learning [33]. We investigate the impact of this learning rule on the self-organisation of the synaptic weights and firing rate responses of the output neurons when all input neurons were modulated by sigmoidal eye-position gain fields. The learning rule was defined by

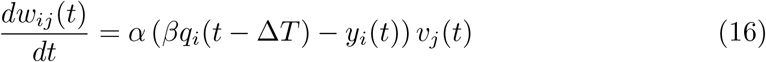

where *y_i_* and *v_j_* were, respectively, the post and presynaptic firing rate values, *α* was the learning rate, *β* was an unconstrained tuning parameter, and *q_i_* was the trace value of the output neuron *i* calculated according to Eq 5 at time (*t* − Δ*T*)*ms*. Eq 16 resembles a form of error-correction learning where the delayed-trace term 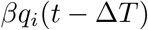 is the target for the current postsynaptic firing rate. The learning rule is biologically plausible in that it utilises only the local activities of the pre- and post-synaptic neurons.

Expanding Eq 16 will result in

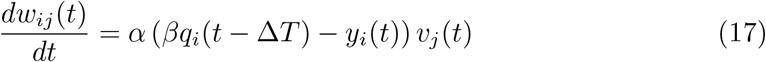

where 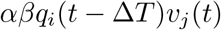 is the *delayed-trace term* of the learning rule. This is the term which contains the tuning parameter ft. The remaining term −*αy_i_*(*t*)*v_j_*(*t*) is minus the learning rate *α* times the product of the post and presynaptic firing rates *y_i_* and *v_j_*, respectively. This term is referred to as the *anti-Hebbian term* of the learning rule.

The behaviour of the learning rule shown in Eq 16 is governed by scaling the parameter *β*. Scaling *β* up could result in the delayed-trace component dominating the learning rule and, therefore, resulting in the same trace-like training behaviour described in previous sections. Similarly, scaling *β* down could make the anti-Hebbian term dominate the behaviour of the learning rule and consequently lead to a qualitative change in the final outcome of the training.

The parameters for the model are given in Table 1. The model was trained according to the description given in section The Visually-Guided Training of the Network. Likewise, in all cases the model was tested on the same visual stimuli. The time delay Δ*T* used to compute the trace value of each output neuron and the parameter *β* were both tuned to optimise the performance of the model at developing head-centred output neurons. The time delay Δ*T* used to compute the trace value of each output neuron *i* was 50*ms*. The parameter *β* was set to 2.2.

Fig 12 shows the firing rate responses of output neuron #168 before and after training, with results shown for four different eye-positions. The miniature scatter plots presented within each of the two subplots **A** and **B** show the reference frame values of all neurons in the output layer. Neuron #168 is indicated in the scatter plots by a red mark. Fig 12**A** shows that prior to training the response of output neuron #168 had no consistent structure in head-centred space across different eye-positions. However, Fig 12**B** shows that after training the output neuron responded maximally to the same head-centred location across all four eye-positions. This neuron has thus learned to respond in a head-centred reference frame.

**Fig 12.**
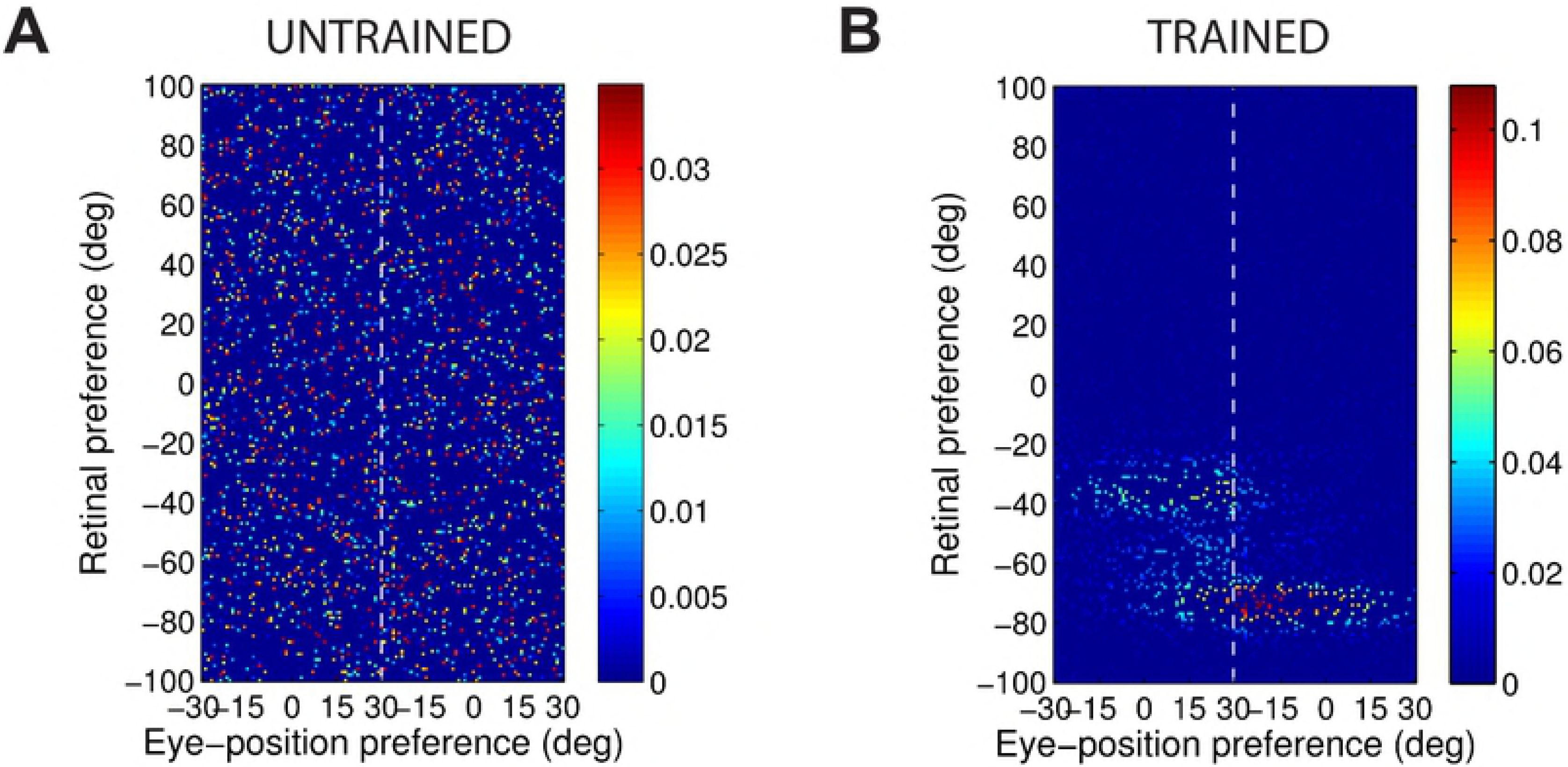
Simulation results showing the firing rate responses of a model incorporating a population of monotonic modulated input neurons trained with the modified learning rule 16: Delayed Postsynaptic Trace with anti-Hebbian Learning. The figure shows the firing rate responses of output neuron #168 before training **(A)** and after training **(B)** during testing for four different eye-positions: −18°, −6°, 6° and 18°. In each subplot, each curve corresponds to a fixed eye-position while a visual target is presented across the same range of head-centred locations. The miniature scatter plot shows the reference frame values of all neurons in the output layer, where each neuron is plotted as a point corresponding to that neuron’s particular combination of head-centredness (ordinate) and eye-centredness (abscissa). The neuron whose firing rate responses have been plotted is shown in the scatter plot by a red mark. After training it is evident that this neuron responds reasonably invariantly to a visual target presented at the same head-centred location regardless of the eye-position.

The change in the synaptic weight structure of the same output neuron #168 due to training is shown in Fig 13. Before training started the afferent synaptic weights were randomly assigned (Fig 13**A**). Fig 13**B** shows that after training, in contrast to the horizontal structure previously obtained with the standard trace learning rule shown in Fig 7**D**, the structure obtained with the modified learning rule 16 is similar to the predicted structure shown in Fig 8 for a head-centred neuron.

**Fig 13.**
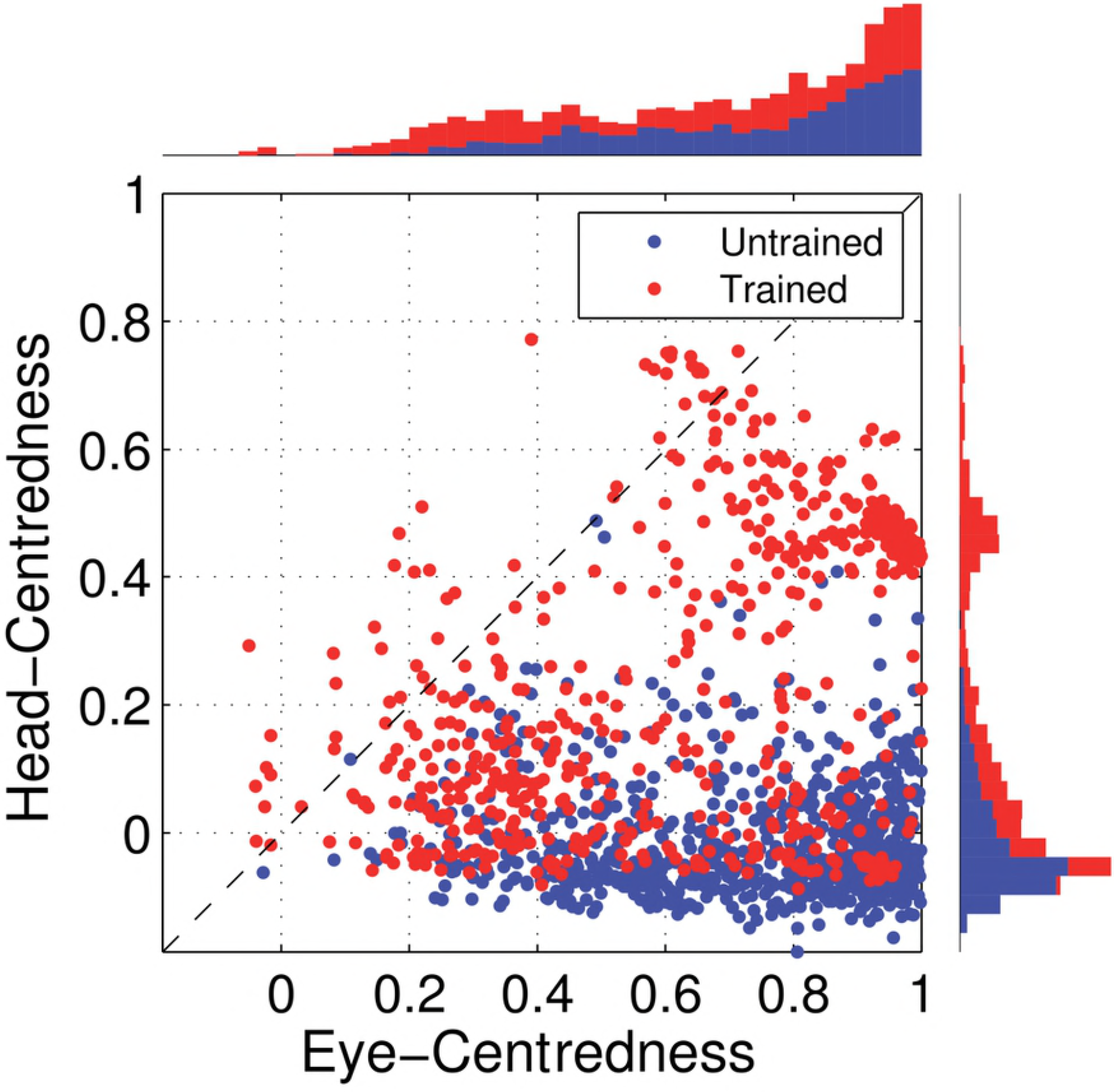
Simulation results showing the strengths of the afferent synapses of a model incorporating a population of sigmoidal modulated input neurons trained with the modified learning rule 16: Delayed Postsynaptic Trace with anti-Hebbian Learning. The figure shows the strengths of the afferent synapses from the input population to output neuron #168 for the untrained **(A)** and trained **(B)** model. The output neuron corresponds to the one plotted in Fig 12. In each plot, the afferent synapses have been arranged topographically by the preference of the input neuron for retinal location *α_i_* and eye-position *β_j_*. The portion of each plot to the left of the white dashed line corresponds to input neurons with positive gain *κ_j_* > 0, whilst the portion of each plot to the right of the white dashed line corresponds to those input neurons with negative gain *κ_j_* < 0. The synaptic weights for this output neuron after training **(B)** have approximately the correct structure for a head-centred neuron as shown in Fig 8.

Fig 14 presents a population analysis of the reference frame response characteristics of the output neurons before and after training. In particular, the scatter plot in Fig 14 shows that before training nearly all of the output neurons had head-centredness values close to 0 and were classified as eye-centred. However, after training the head-centredness values of many output neurons had dramatically increased, with quite a number of these neurons now classed as head-centred. Comparing the output population analysis of the model trained with the modified learning rule 16 shown in Fig 14 with the performance of the model trained with the standard trace learning rule (6) shown in Fig 5 confirms that the new modified learning rule 16 is far more efficacious at driving the development of head-centred output neurons when the input neurons are modulated by a sigmoidal (monotonic) function of eye-position than the standard trace learning rule (6) originally investigated by [1].

**Fig 14.**
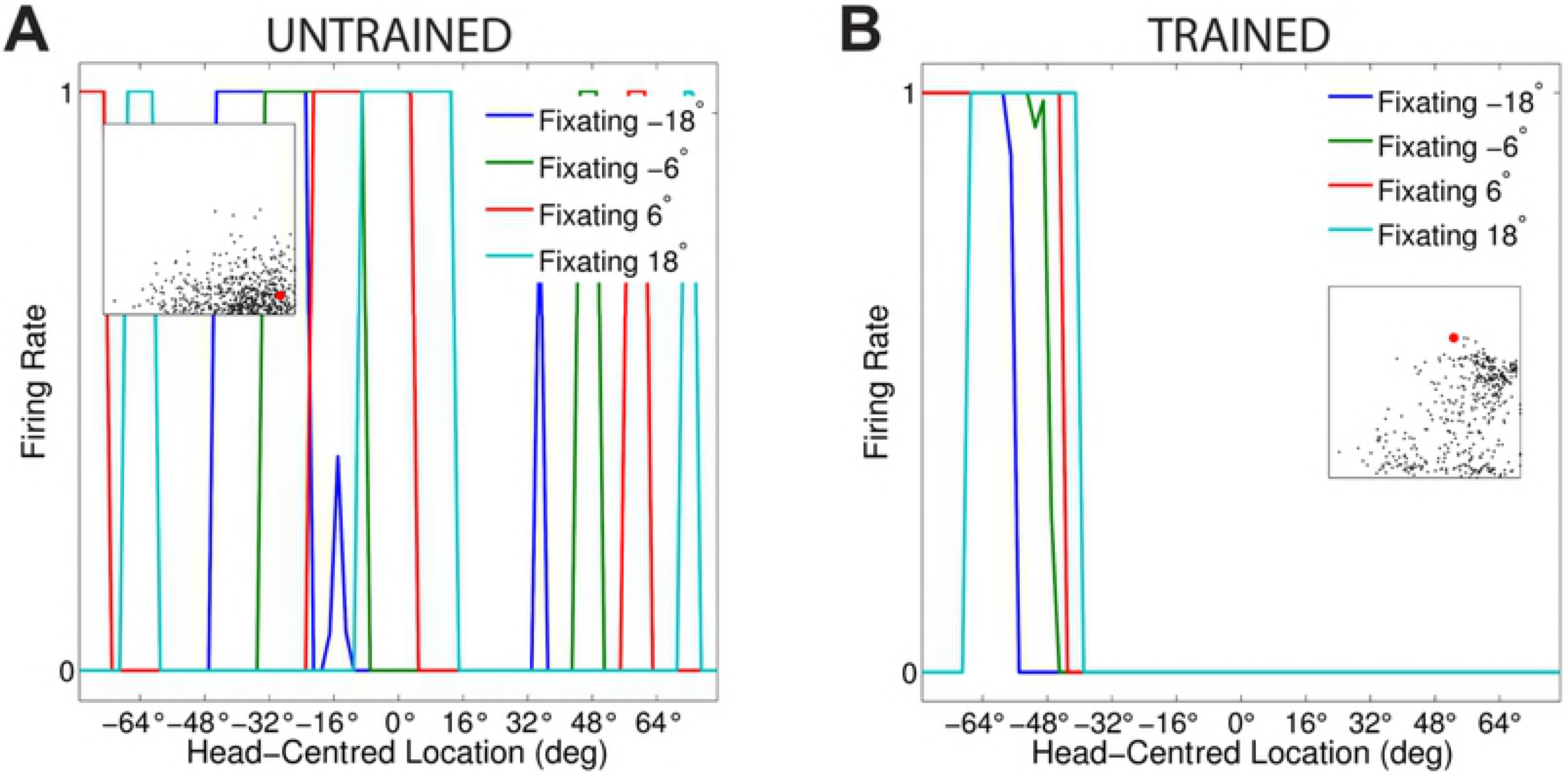
Simulation results showing the output reference frame response characteristics of a model incorporating a population of sigmoidal modulated input neurons trained with the modified learning rule 16: Delayed Postsynaptic Trace with anti-Hebbian Learning. The scatter plot shows the reference frame response characteristics of all output neurons before and after training. Each neuron is represented as a point corresponding to its combination of eye-centredness (abscissa) and head-centredness (ordinate) values. Data points for the untrained model are plotted in blue and data points for the trained model are shown in red. The dashed diagonal line with positive unity slope separates those neurons which are classified as head-centred (above the line) from those that are classified as eye-centred (below the line). It can be seen that many of the output neurons have developed head-centred output responses after training.

In summary, these results showed that training the model with the modified learning rule 16 refined the response characteristics of many output neurons to be more compatible with a head-centred frame of reference, even when all of the input neurons had monotonic eye-position gain modulation.

### Modified Learning Rule: Delayed Postsynaptic Firing Rate with Anti-Hebbian Learning

This section presents simulation results showing the performance of a learning rule which incorporates a delayed postsynaptic firing rate with anti-Hebbian learning [25]. The same training procedure of previous simulations was used in the simulations presented in this section. The learning rule is defined by

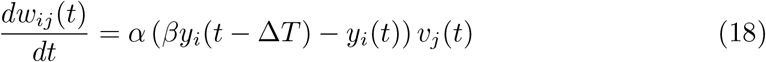

where *y_i_* was the firing rate of output neuron *i*. In this case no postsynaptic trace value *q_i_* is used to update the synaptic weights. The tuning parameter *β* works as described in section Modified Learning Rule: Delayed Postsynaptic Trace with Anti-Hebbian Learning for the anti-Hebbian learning rule with delayed trace (Eq 16). The time delay Δ*T* in the firing rate of each output neuron and the parameter *β* were both tuned to optimise the performance of the model in driving the development of head-centred output neurons. The value of the time delay Δ*T* was set to 500*ms* and *β* was set to 2.4. Simulation parameters for the model are shown in Table 1.

Fig 15 shows the firing rate responses of output neuron #876 before training (Fig 15**A**) and after training (Fig 15**B**). Fig 15**B** shows that after training output neuron #876 responded to the same head-centred location across different eye-positions. This was not the case for the same output neuron before training (Fig 15**A**). Thus, it is evident that during training the neuron has learned to respond in a head-centred reference frame.

**Fig 15.**
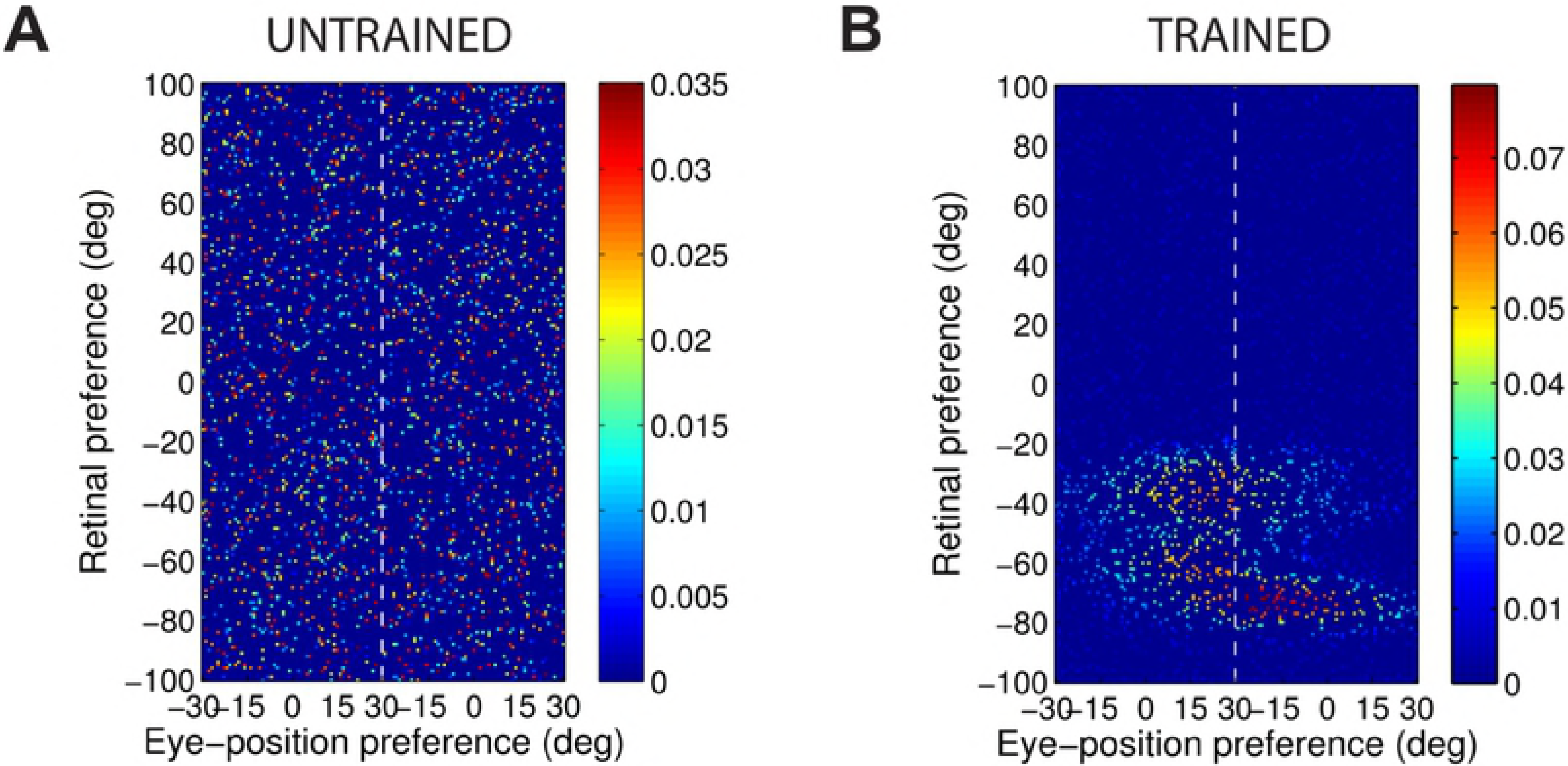
Simulation results showing the firing rate responses of a model incorporating a population of monotonic modulated input neurons trained with the modified learning rule 18: Delayed Postsynaptic Firing Rate with anti-Hebbian Learning. The figure shows the firing rate responses of output neuron #876 before training **(A)** and after training **(B)** during testing for four different eye-positions: −18°, −6°, 6° and 18°. Conventions as for Fig 12. The comparison of subplot **(A)** and subplot **(B)** shows that the output neuron learned to respond to a specific head-centred location regardless of the eye-position after training.

Fig 16 shows how the synaptic weight structure of the same output neuron #876 changed due to training the model with learning rule 18. The final weight structure after training shown in Fig 16B resembles the predicted weight structure shown in Fig 8 for head-centred neurons.

**Fig 16.**
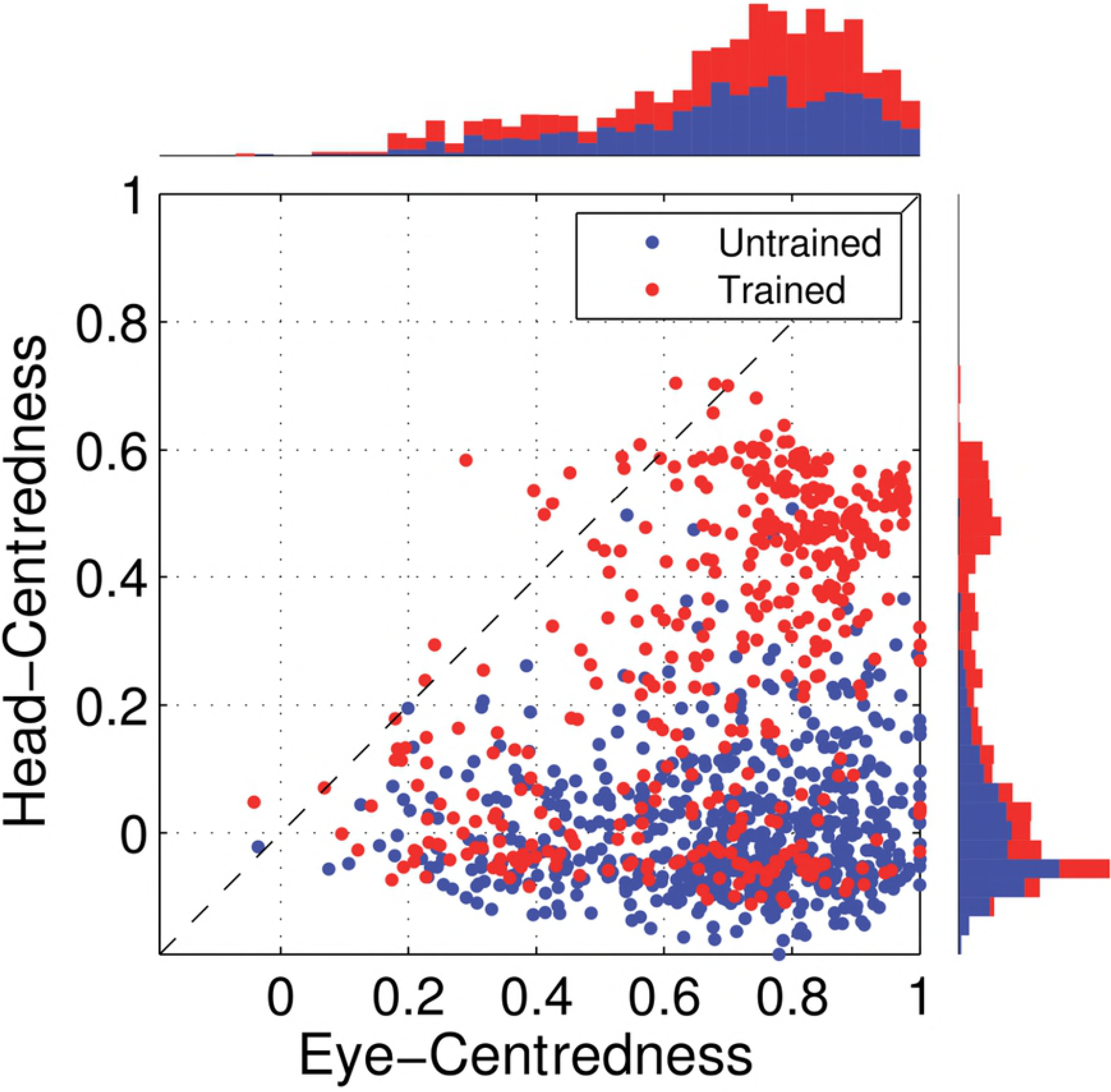
Simulation results showing the strengths of the afferent synapses of a model incorporating a population of sigmoidal modulated input neurons trained with the modified learning rule 18: Delayed Postsynaptic Firing Rate with anti-Hebbian Learning. The figure shows the strengths of the afferent synapses from the input population to output neuron #876 for the untrained **(A)** and trained **(B)** model. The output neuron corresponds to the one plotted in Fig 15. Conventions as for Fig 13. The synaptic weight structure for this output neuron after training shown in plot **(B)** has approximately the correct profile for a head-centred neuron (Fig 8).

Fig 17 shows the eye-centredness and head-centredness values of output neurons in the untrained and in the trained model. It is clear that almost none of the output neurons in the untrained model had values of head-centredness higher than eye-centredness. However, after training the head-centredness values of many output neurons increased substantially, with a number of such neurons now having greater head-centredness than eye-centredness values. Such neurons are, therefore, classified as head-centred neurons (section Analysis of Network Performance). Comparing the output population analysis of the model trained with the modified learning rule 18 shown in Fig 17 with the analysis of the model trained with the standard trace learning rule (6) shown in Fig 5 demonstrates that the new modified learning rule 18 is also more efficacious at producing head-centred output neurons when the input neurons are modulated by a sigmoidal function of eye-position than the standard trace learning rule (6) previously implemented by [1].

**Fig 17.**
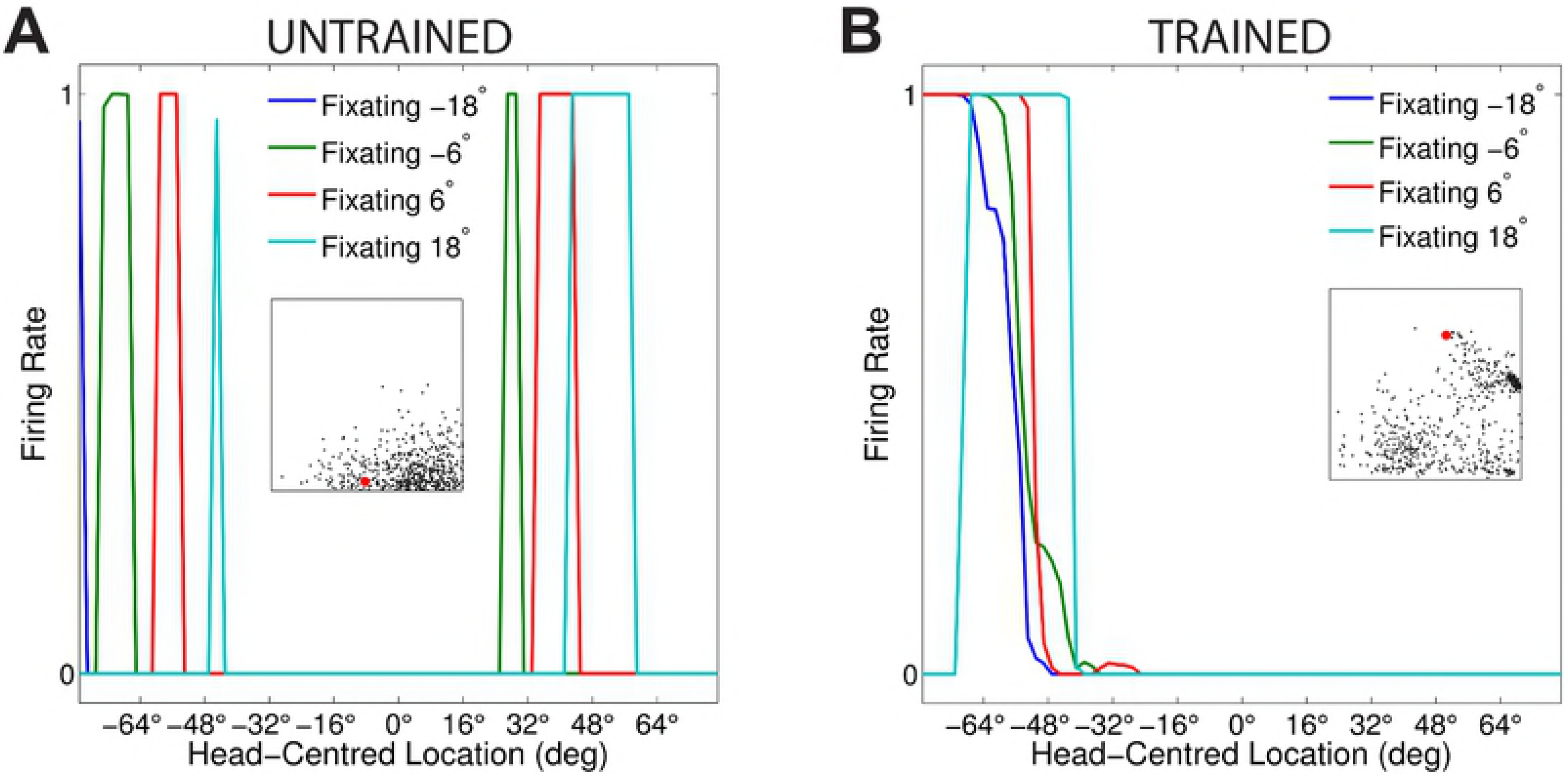
Simulation results showing the output reference frame response characteristics of a model incorporating a population of sigmoidal modulated input neurons trained with the modified learning rule 18: Delayed Postsynaptic Firing Rate with anti-Hebbian Learning. The scatter plot shows the reference frame response characteristics of all output neurons before and after training. Conventions as for Fig 14. It is evident that training had the effect of increasing the head-centredness values of most output neurons. Indeed many more head-centred output neurons are present in the trained model than in the untrained model.

In summary, these results showed that training the model with the modified learning rule 18 resulted in the development of head-centred output neurons, even when the whole input population had sigmoidal eye-position gain modulation. However, the comparison of Fig 17 and Fig 14 shows that learning rule 16, which incorporated a delayed postsynaptic trace *q_i_*(*t* − Δ*T*), is in fact more efficacious at driving the development of head-centred output neurons than learning rule 18, which incorporated the delayed firing rate *y_i_*(*t* − Δ*T*). Thus, the incorporation of the trace value *q_i_*(*t* − Δ*T*) enhances the ability of the learning rule to perform temporal binding of input patterns corresponding to the same head-centred location.

### Modified Learning Rule: Current Postsynaptic Trace with Anti-Hebbian Learning

This section presents simulation results showing the performance of a modified learning rule which incorporated the current postsynaptic trace with anti-Hebbian learning. Thus, this learning rule does not use an explicit time delay Δ*T*. The learning rule was given by

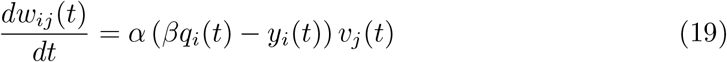

where *α* was the learning rate, *β* was the tuning parameter, *q_i_* was the trace value of the output neuron *i*, and *y_i_* and *v_j_* were the post- and pre-synaptic firing rate values, respectively. What distinguishes the learning rule in Eq 19 from the learning rule in Eq 16 is the use of the trace value calculated at the same time *t* when the synaptic weight is updated.

Table 1 gives the parameters for the model. The value of the tuning parameter *β* was set to 2.2 for optimal performance.

Fig 18 shows the firing rate responses of output neuron #328 for the untrained and trained models. Specifically, Fig 18**A** shows that prior to training the response of output neuron #328 had no consistent structure in head-centred space across different eye-positions. However, Fig 18**B** shows that after training the same output neuron reponded maximally to the same head-centred location across all four eye-positions. Thus, after training, neuron #328 responded in a head-centred frame of reference.

**Fig 18.**
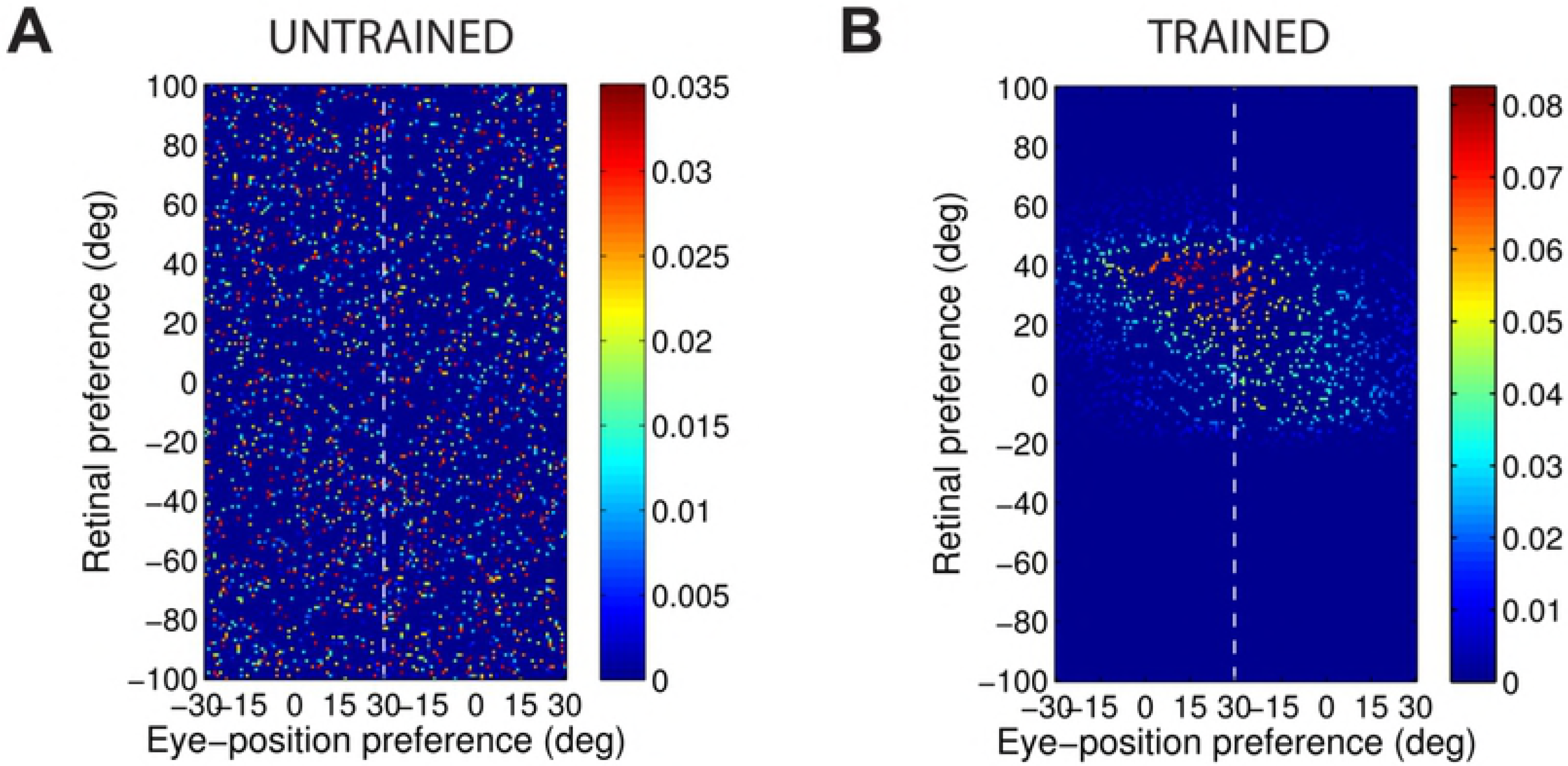
Simulation results showing the firing rate responses of a model incorporating a population of monotonic modulated input neurons trained with the modified learning rule 19: Current Postsynaptic Trace with anti-Hebbian Learning. The figure shows the firing rate responses of output neuron #328 before training **(A)** and after training **(B)** during testing for four different eye-positions: −18°, −6°, 6° and 18°. Conventions as for Fig 12. The comparison of subplot **(A)** and subplot **(B)** shows that the output neuron learned to respond to a specific head-centred location regardless of the eye-position after training.

The change in the synaptic weight structure of the same output neuron #328 due to 975 training is shown in Fig 19. Before training the afferent synaptic weights were randomly 976 assigned (Fig 19**A**). After training, however, the synaptic weight structure of the same 977 output neuron (Fig 19**B**) is similar to the predicted weight structure for head-centred neurons shown in Fig 8.

**Fig 19.**
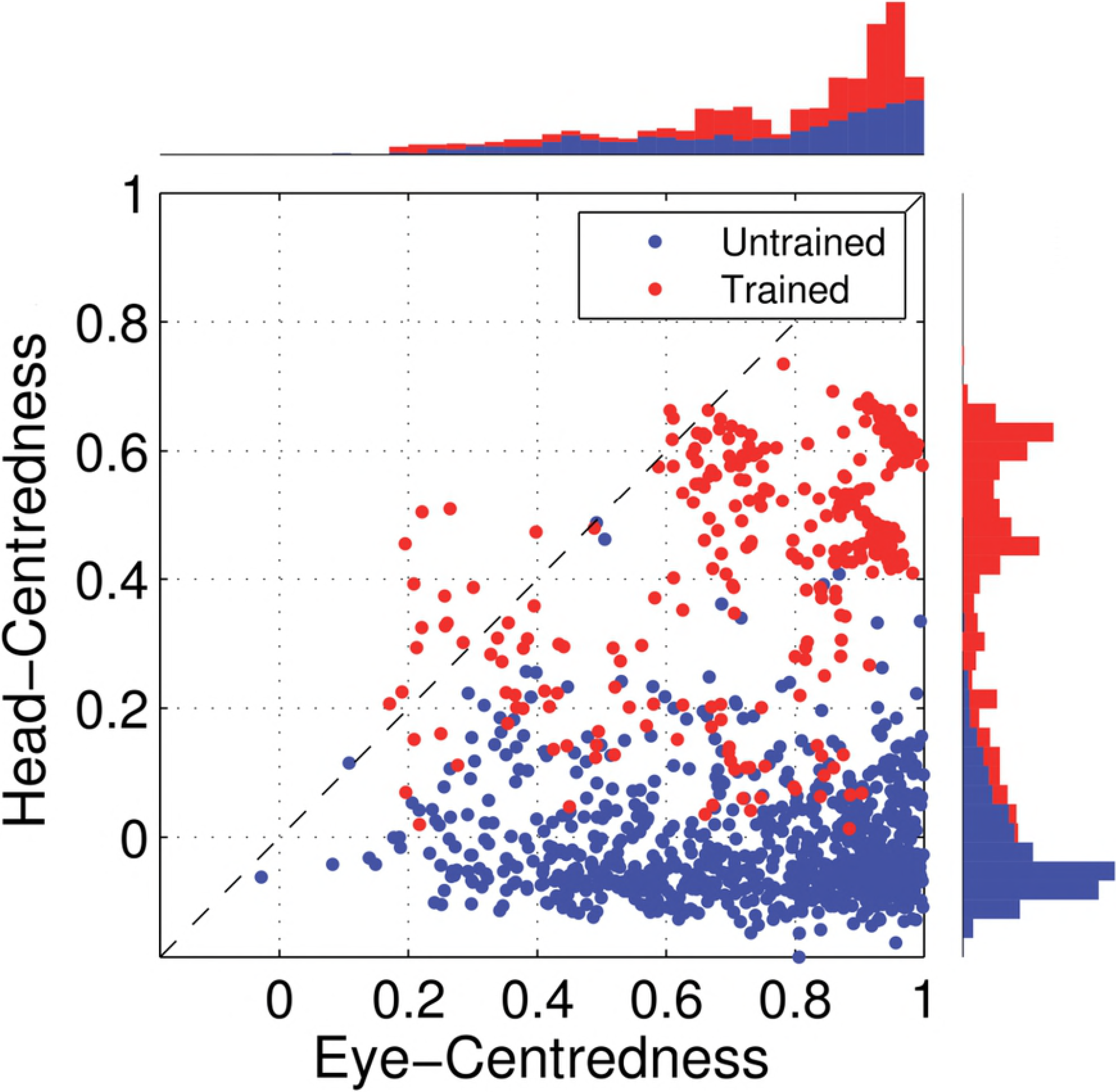
Simulation results showing the strengths of the afferent synapses of a model incorporating a population of sigmoidal modulated input neurons trained with the modified learning rule 19: Current Postsynaptic Trace with anti-Hebbian Learning. The figure shows the strengths of the afferent synapses from the input population to output neuron #328 for the untrained **(A)** and trained **(B)** model. The output neuron corresponds to the one plotted in Fig 18. Conventions as for Fig 13. The synaptic weight structure for this output neuron after training shown in plot **(B)** has the correct kind of profile for a head-centred neuron (Fig 8).

Fig 20 shows the eye-centredness and head-centredness values of output neurons in the untrained and in the trained model. In particular, Fig 20 shows that training had the effect of increasing the head-centredness value for a large proportion of output neurons. Furrthermore, while almost no output neurons were classified as head-centred before training, a significant number of output neurons were classified as head-centred after training. A comparison of the output population analysis of the model trained with the modified learning rule 19 shown in Fig 20 with the performance of the model trained with the standard trace learning rule (6) presented in Fig 5 shows that the modified learning rule 19 is also significantly more capable of producing head-centred output neurons than the standard trace learning rule (6) when all of the input neurons have sigmoidal gain modulation by eye-position.

**Fig 20.**
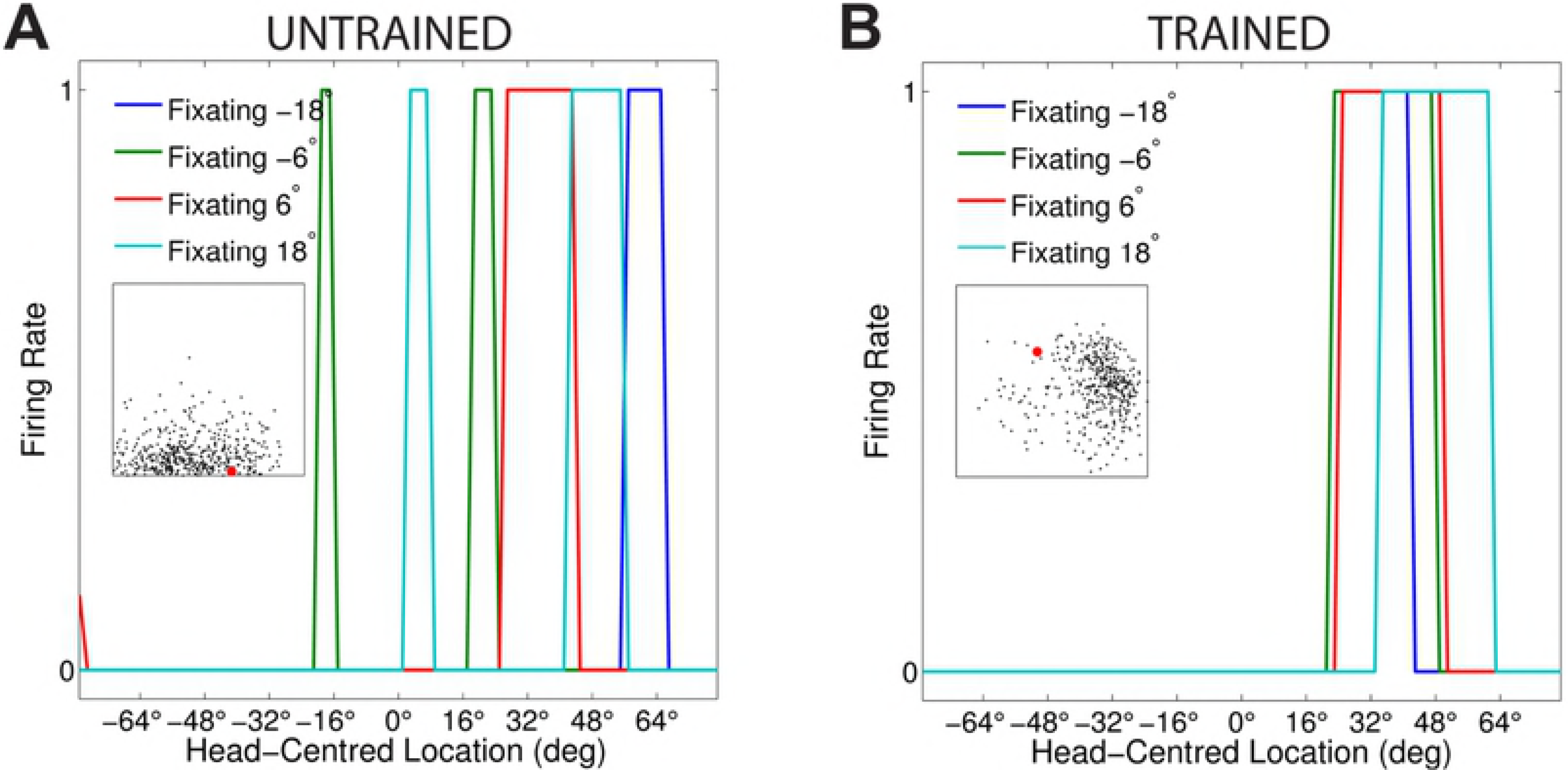
Simulation results showing the output reference frame response characteristics of a model incorporating a population of sigmoidal modulated input neurons trained with the modified learning rule 19: Current Postsynaptic Trace with anti-Hebbian Learning. The scatter plot shows the reference frame response characteristics of all output neurons before and after training. Conventions as for Fig 14. It can be seen that training increased the head-centredness values of most output neurons, with quite a number of head-centred output neurons present in the trained model.

In summary, these results showed that training the model with learning rule 19 drives the development of head-centred output neurons even when the whole input population had sigmoidal eye-position gain modulation. The comparison of Fig 20 with Fig 14 shows that learning rule 16, which incorporated a delayed postsynaptic trace *q_i_*(*t* − Δ*T*), is more effective at producing head-centred output neurons than learning rule 19, which incorporated the current trace *q_i_*(*t*) calculated at the same time *t* when the weights are updated.

### Modified Learning Rule: Delayed Postsynaptic Trace Learning Rule

The simulation results presented in section Modified Learning Rule: Delayed Postsynaptic Trace with Anti-Hebbian Learning, section Modified Learning Rule: Delayed Postsynaptic Firing Rate with Anti-Hebbian Learning and section Modified Learning Rule: Current Postsynaptic Trace with Anti-Hebbian Learning showed, respectively, that training the model with learning rule 16, learning rule 18 and learning rule 19 successfully self-organised head-centred output responses and increased the head-centredness value of a large proportion of output neurons when all input neurons had sigmoidal eye-position gain modulation. Importantly, section Self-organisation with peaked and monotonic gain fields showed this was not the case for the standard trace learning rule (6). Training the model with the standard trace learning rule successfully self-organised head-centred output neurons when all of the input neurons had peaked eye-position gain modulation, but failed when the modulation of the input population was altered from peaked to sigmoidal. Indeed, the simple introduction of a small proportion (e.g. with *p* > 0.2) of input neurons with sigmoidal eye-position gain modulation was enough to undermine the self-organisation of head-centred output responses (section Standard Trace Learning Rule with Mixed Peaked and Sigmoidal eye-position Modulation of Input Neurons).

The modified learning rules introduced in section Modified Learning Rule: Delayed Postsynaptic Trace with Anti-Hebbian Learning, section Modified Learning Rule: Delayed Postsynaptic Firing Rate with Anti-Hebbian Learning and in section Modified Learning Rule: Current Postsynaptic Trace with Anti-Hebbian Learning all had an anti-hebbian term as their common component. Out of these learning rules, the best performance was observed with learning rule 16, which incorporated a postsynaptic delayed-trace term. In this case, an interesting question is whether the superior efficacy of this learning rule in driving the development of head-centred output responses was primarily due to the anti-hebbian term or the postsynaptic delayed-trace term. In particular, is an anti-hebbian term actually needed in this learning rule for the self-organisation of head-centred responses in the presence of input neurons with sigmoidal eye-position gain modulation, or could head-centred output responses develop using a learning rule that only incorporated a postsynaptic delayed-trace term?

This section addresses the above questions by investigating the performance of the model with the following learning rule

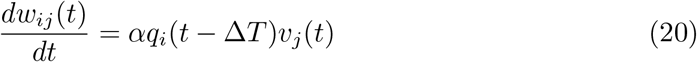

where *α* was the learning rate, *q_i_* was the trace value of the output neuron *i* and *V_j_* was the firing rate of the input neuron *j*. In Eq 20 the trace value *q_i_* is calculated at time (*t* − Δ*T*)*ms*. This learning rule has no anti-hebbian term, and relies purely on the postsynaptic delayed-trace term *q_i_*(*t* − Δ*T*) to drive the development of head-centred output neurons.

The simulation parameters for the model are given in Table 1. The time delay Δ*T* used to compute the trace value of each output neuron *i* was set to 30*ms* for optimal learning performance.

Fig 21 shows how training changed the firing rate responses of output neuron #223. Prior to training the response of output neuron #223 had no consistent structure in head-centred space across different eye-positions (Fig 21 **A**), whilst after training the same output neuron responded maximally to the same head-centred location across all four eye-positions (Fig 21**B**). Thus, neuron #223 learned to respond in a head-centred reference frame after training.

**Fig 21.**
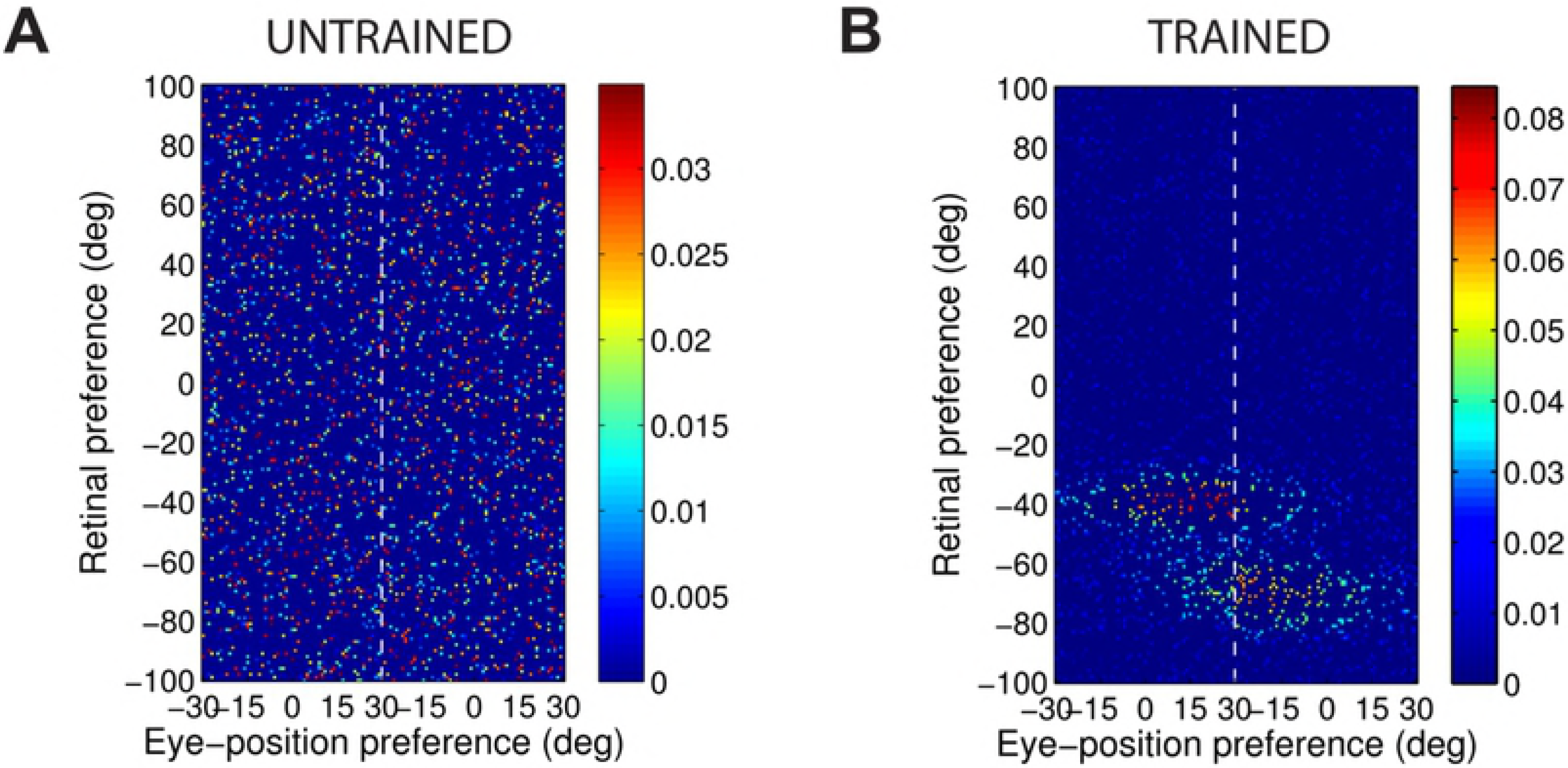
Simulation results showing the firing rate responses of a model incorporating a population of monotonic modulated input neurons trained with the modified learning rule 20: Delayed Postsynaptic Trace Learning Rule. The figure shows the firing rate responses of output neuron #223 before training **(A)** and after training **(B)** during testing for four different eye-positions: −18°, −6°, 6° and 18°. Conventions as for Fig 12. The comparison of subplot **(A)** and subplot **(B)** shows that the output neuron learned to respond to a specific head-centred location regardless of the eye-position after training. Output neuron #223 is not shown in the scatter plot of subplot **(A)** because this neuron did not respond for every eye-position before training and was therefore excluded from further analysis (section Analysis of Network Performance). However, subplot **(B)** shows that the same output neuron learned to respond to a specific head-centred location regardless of the eye-position after training.

Fig 22 shows how the synaptic weight structure of the same output neuron #223 plotted in Fig 21 changed due to training the model with learning rule 20. It is clear that the final synaptic weight structure after training (Fig 22**B**) resembles the predicted weight structure for head-centred neurons (Fig 8), even though this was not the case before training (Fig 22**A**).

**Fig 22.**
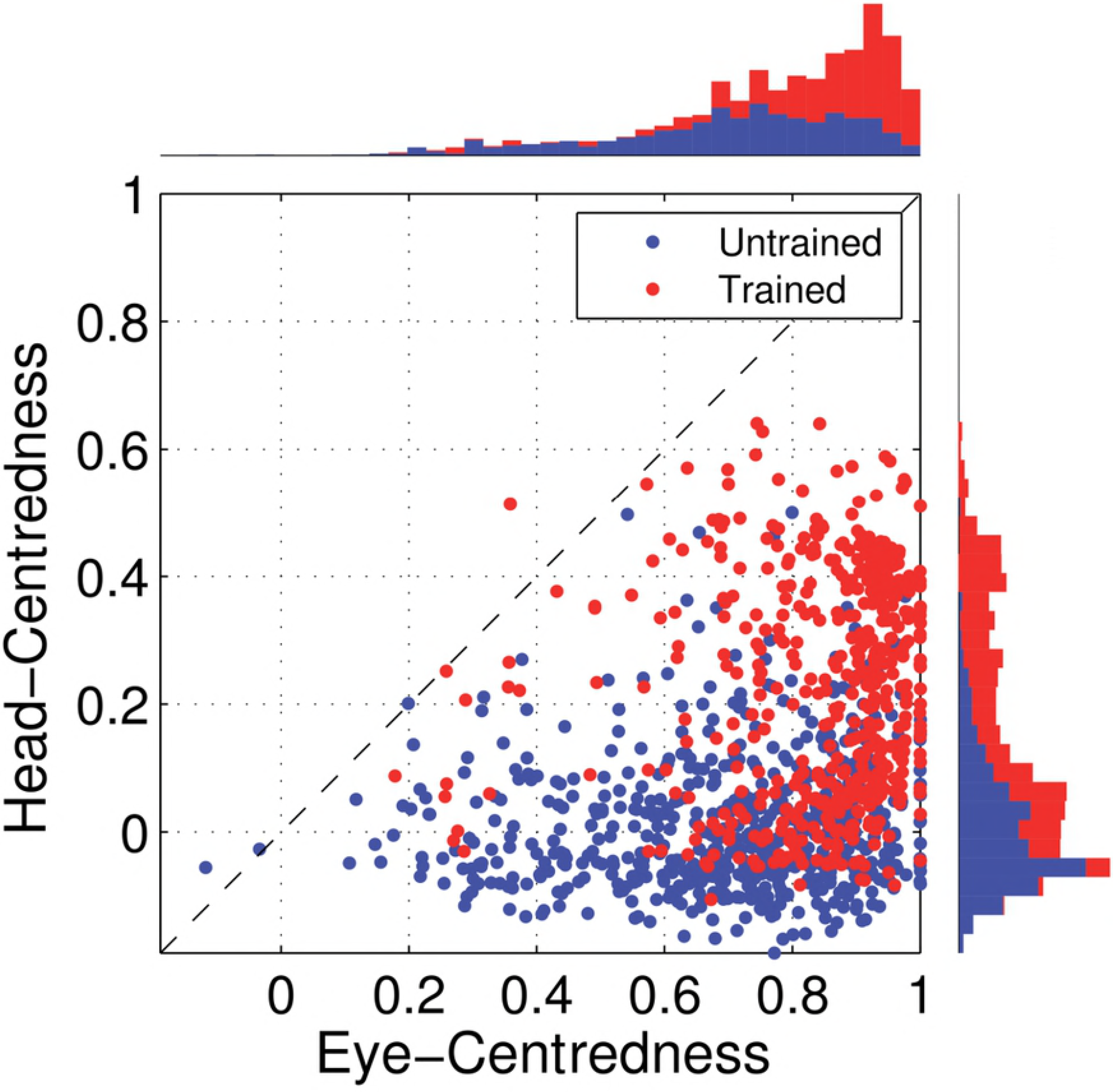
Simulation results showing the strengths of the afferent synapses of a model incorporating a population of sigmoidal modulated input neurons trained with the modified learning rule 20: Delayed Postsynaptic Trace Learning Rule. The figure shows the strengths of the afferent synapses from the input population to output neuron #223 for the untrained **(A)** and trained **(B)** model. The output neuron corresponds to the one plotted in Fig 21. Conventions as for Fig 13. The synaptic weight structure for this output neuron after training shown in plot **(B)** has approximately the correct profile for a head-centred neuron (Fig 8).

Fig 23 shows the eye-centredness and head-centredness values of all of the output neurons in the untrained model and in the trained model. After training, the head-centredness values of many output neurons had increased substantially. However, there was only a single output neuron, which was neuron #223, with a greater head-centredness value than eye-centreredness, and which was consequently classed as responding in a head-centred reference frame. Therefore, in the simulations presented in this paper, the anti-hebbian term in learning rules (16), (18) and (19) appears to play an important and essential role in producing relatively large numbers of head-centred output neurons when all of the input neurons have sigmoidal eye-position gain fields.

**Fig 23.**
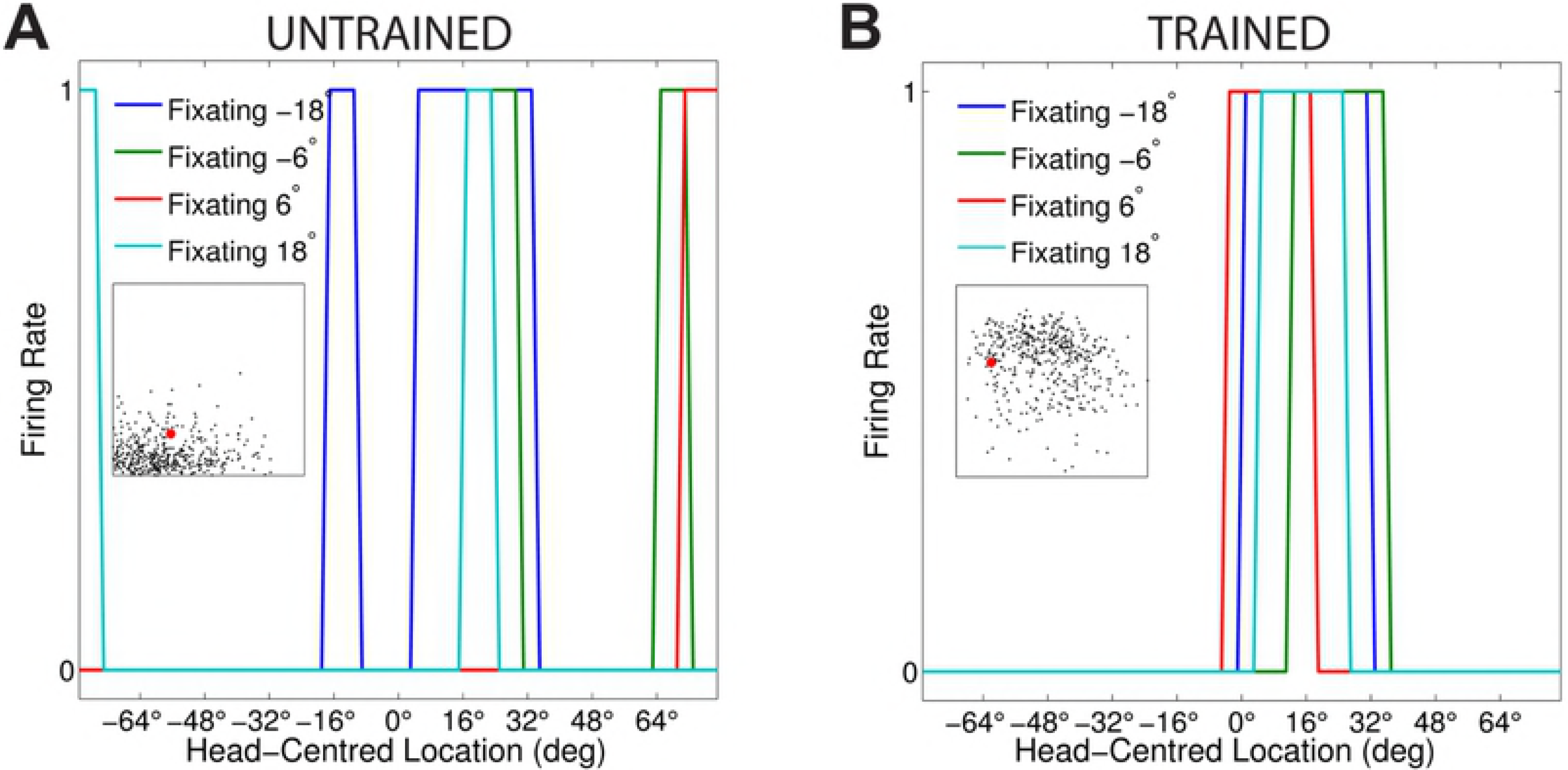
Simulation results showing the output reference frame response characteristics of a model incorporating a population of sigmoidal modulated input neurons trained with the modified learning rule 20: Delayed Postsynaptic Trace Learning Rule. The scatter plot shows the reference frame response characteristics of all output neurons before and after training. Conventions as for Fig 14. It is evident that training had the effect of increasing the head-centredness values of most output neurons. Although, there is only a single head-centred output neuron present in the trained model.

### Delayed Postsynaptic Trace Learning Rule with Mixed Peaked and Sigmoidal eye-position Modulation of Input Neurons

In the previous section, it was found that an anti-hebbian term was needed in the learning rule in order to produce relatively large numbers of head-centred output neurons if the entire population of input neurons had sigmoidal gain fields. However, experimental studies have demonstrated that the primate cortex contains a mixed population of visual neurons with either peaked or monotonic eye-position gain fields [7–9]. This raises the question of whether learning rule (20), which relies solely on a postsynaptic delayed-trace term without any anti-hebbian term, could produce a much larger number of head-centred output neurons if the input population had a 50:50 mix of peaked and sigmoidal eye-position gain fields.

This section presents simulation results with the modified learning rule (20) when there is a 50:50 mixture of peaked and monotonic gain fields in the visual input population. Can the model produce a large number of head-centred output neurons under such conditions without an anti-hebbian term in the learning rule? This is still a potentially challenging task for the model because in section Standard Trace Learning Rule with Mixed Peaked and Sigmoidal eye-position Modulation of Input Neurons it was shown that the standard trace learning rule (6), which lacks an anti-hebbian term, failed to produce significant numbers of head-centred output neurons when the proportion of sigmoidal gain modulated input neurons rose above just 20% of the overall input population. The model parameters for the simulations presented in this section are given in Table 1.

Fig 24 shows how training changed the firing rate responses of output neuron #281. In particular, Fig 24**B** shows that after training output neuron #281 responded maximally to the same head-centred location across all four eye-positions, although this was not the case prior to training (Fig 24**A**). Hence neuron #281 responded in a head-centred frame of reference after training.

**Fig 24.**
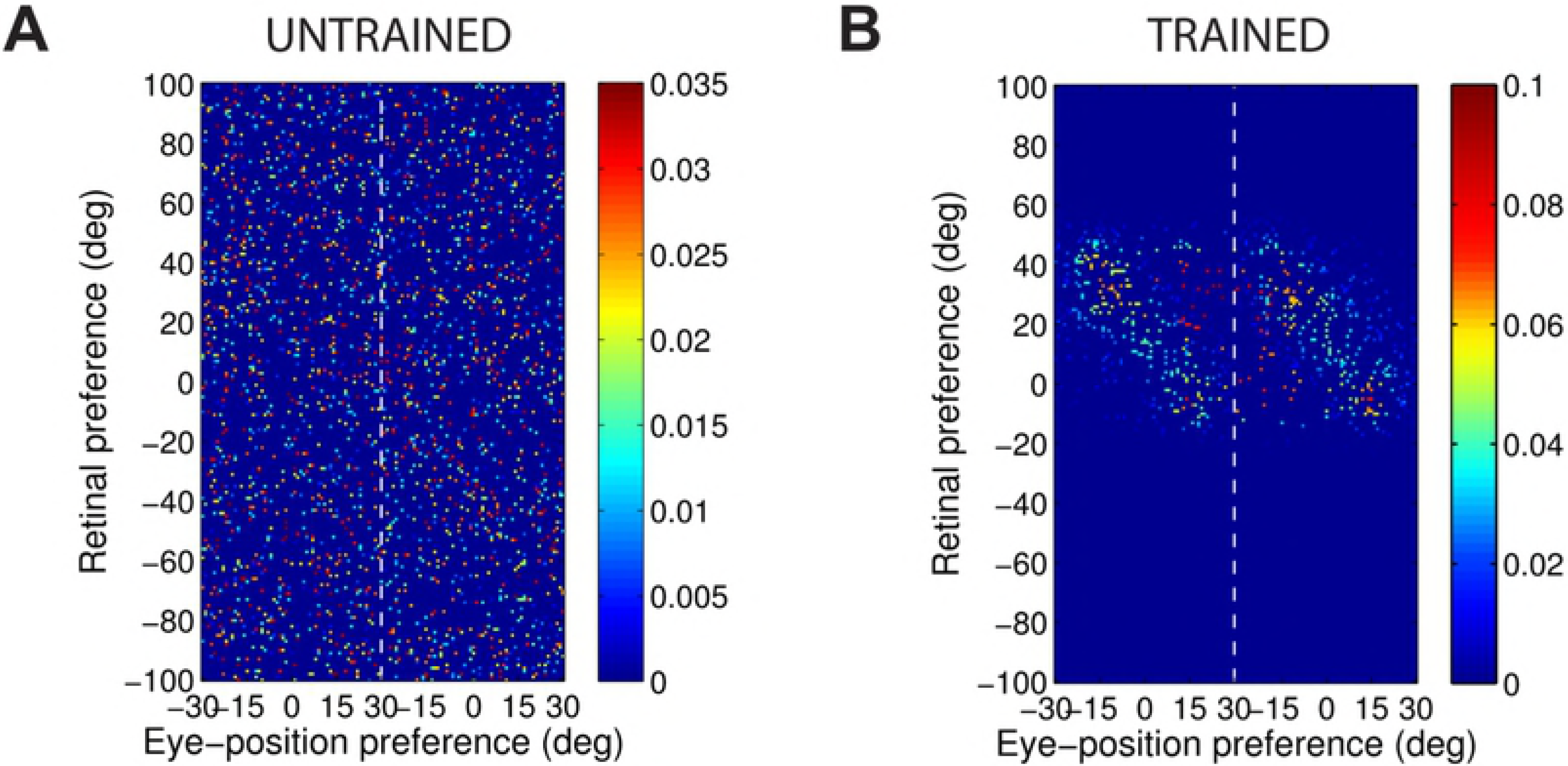
Simulation results of a model incorporating a mixed population of peaked and sigmoidal modulated input neurons, with sigmoidal modulation rate p set to 0.5, trained with the modified learning rule 20: Delayed Postsynaptic Trace Learning Rule. The figure shows the firing rate responses of output neuron #281 before training **(A)** and after training **(B)** during testing for four different eye-positions: −18°, −6°, 6° and 18°. Conventions as for Fig 12. Plot **(B)** shows that the same output neuron learned to respond to a specific head-centred location regardless of the eye-position after training.

Fig 25 shows how the synaptic weight structure of the same output neuron #281 plotted in Fig 24 changed due to training. Fig 25**B** shows that after training, the synaptic weight structure of the output neuron was clearly similar to the predicted weight structure for head-centred neurons shown in Fig 8. The synaptic weight structure before training reflected the randomly assigned connection weights (Fig 25**A**).

**Fig 25.**
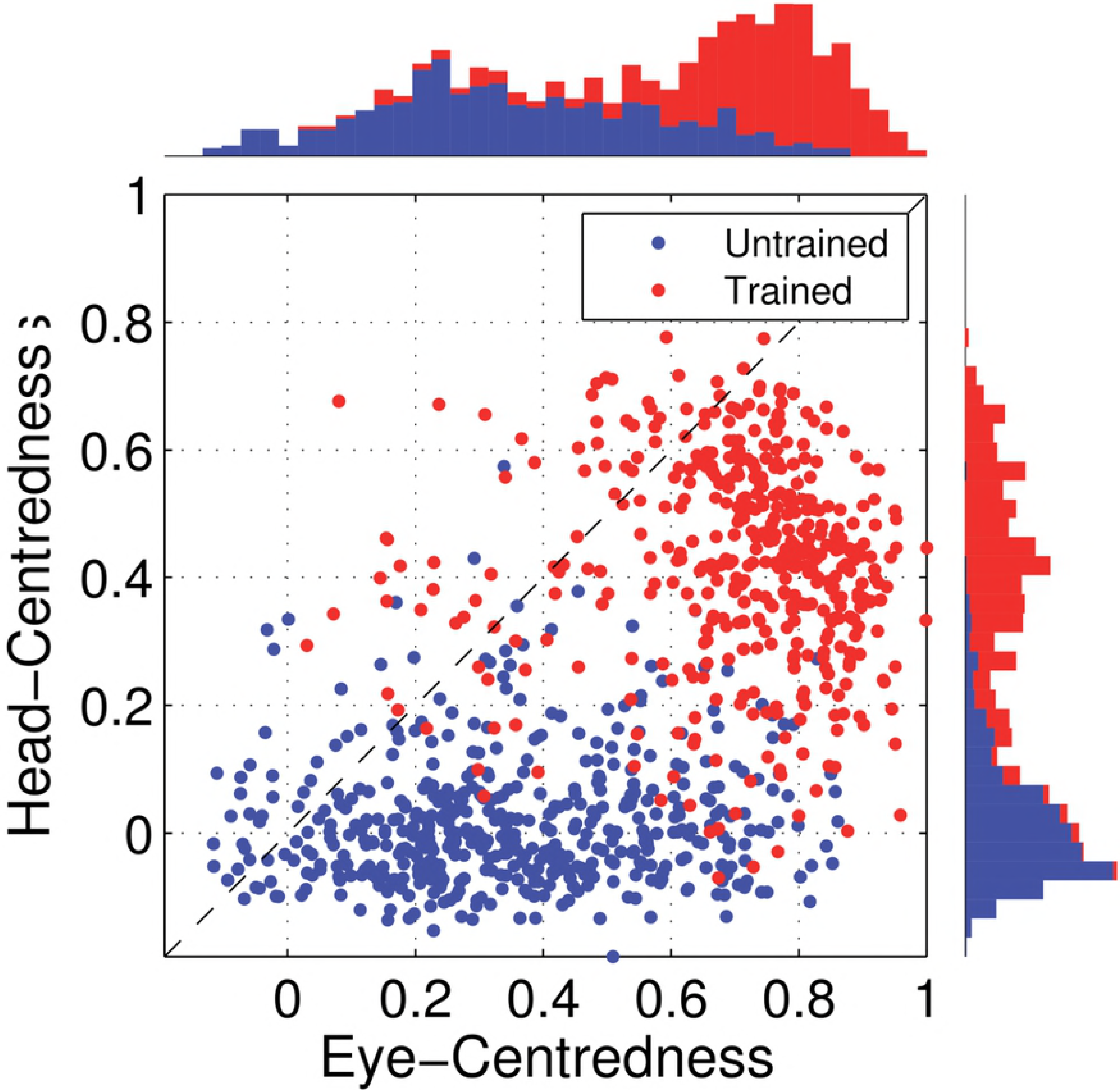
Simulation results of a model incorporating a mixed population of peaked and sigmoidal modulated input neurons, with sigmoidal modulation rate *p* set to 0.5, trained with the modified learning rule 20: Delayed Postsynaptic Trace Learning Rule. The figure shows the strengths of the afferent synapses from the input population to output neuron #281 for the untrained **(A)** and trained **(B)** model. The output neuron corresponds to the one plotted in Fig 24. Conventions as for Fig 13. The synaptic weight structure for this output neuron after training shown in plot **(B)** has the correct kind of profile for a head-centred neuron (Fig 8).

Fig 26 shows the eye-centredness and head-centredness values of output neurons in the untrained and in the trained model. Fig 26 shows that training the model with learning rule 20, where the input population contained a 50:50 mix of neurons with peaked or sigmoidal gain modulation, had the effect of increasing the head-centredness value of most output neurons. Moreover, a large proportion of output neurons in the trained model are head-centred (i.e. with a head-centredness value greater than eye-centredness). Indeed, more head-centred output neurons were observed in the trained model in this section than in any of the trained models presented in previous section Modified Learning Rule: Delayed Postsynaptic Trace with Anti-Hebbian Learning, section Modified Learning Rule: Delayed Postsynaptic Firing Rate with Anti-Hebbian Learning and section Modified Learning Rule: Current Postsynaptic Trace with Anti-Hebbian Learning.

**Fig 26.**
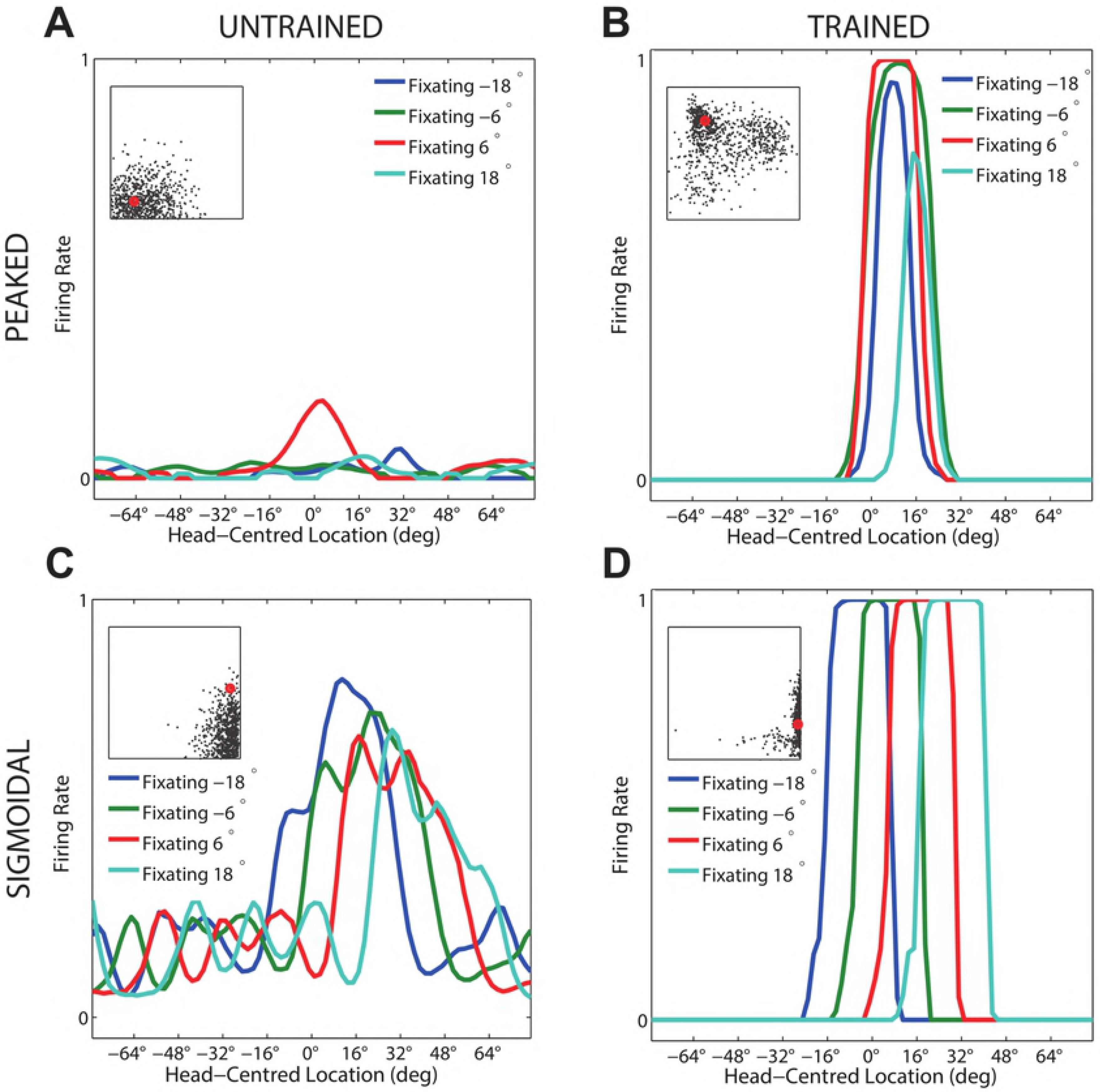
Simulation results of a model incorporating a mixed population of peaked and sigmoidal modulated input neurons, with sigmoidal modulation rate *p* set to 0.5, trained with the modified learning rule 20: Delayed Postsynaptic Trace Learning Rule. The scatter plot shows the reference frame response characteristics of all output neurons before and after training. Conventions as for Fig 14. It is evident that training had the effect of increasing the head-centredness values of most output neurons, with many more head-centred output neurons present in the trained model.

In summary, these results demonstrated that it was possible for the model to produce large numbers of head-centred output neurons with a learning rule that relied purely on a postsynaptic delayed-trace term without any anti-hebbian term if the input population contained a 50:50 mix of neurons that were modulated by either peaked or sigmoidal gain fields. Such a mixture of different forms of eye-position gain modulation is biologically compatible with experimental findings [7,8].

## Discussion

The majority of previously published models of coordinate transformation from eye-centred to head-centred visual representations have relied on some form of supervised error-correction learning, in which an explicit supervisory signal is used to specify the desired head-centred output responses during training [14–16]. The availability of such a supervisory training signal makes the self-organisation of these models robust even with monotonic (e.g. planar or sigmoidal) eye-position gain modulated input neurons. However, an immediate problem with these kinds of models is explaining exactly where such a supervisory training signal might originate from in the brain. Another major problem for models that employ a backpropagation of error supervised learning, such as the classic model of [14], is that this model architecture is entirely biologically implausible [18]. In particular, there is no plausible explanation for how the error terms needed to adjust the afferent synaptic weights in the intermediate layer of the model could be generated and implemented in the brain. One consequence of this is that the backpropagation of error learning procedure can result in individual neurons making both excitatory and inhibitory synaptic connections on different postsynaptic neurons. Although not shown here, we have verified this through replicating the model simulations of [14] and [16]. This feature of backpropagation of error models violates an accepted principle of cortical architecture, sometimes referred to as Dale’s Law, that an individual neuron must have the same kind of excitatory or inhibitory effect at all of its synaptic connections with other neurons [19]. This failure of backpropagation of error models even undermines the potential relevance of their trained synaptic architecture to understanding how coordinate transformations to head-centred visual representations might be implemented in the brain. [15] also used error-correction learning to develop head-centred output representations. Their learning scheme employed a simpler global error signal for the entire output population, which might conceivably be implemented by some form of neuromodulator release. However, the implementation of error correction learning in their model still required complex architectural features that have not been identified in cortex.

To remedy the lack of biological plausibility in previously published models that use supervised learning to drive the development of visual neurons that respond in a head-centred reference frame, [1] demonstrated a model that self-organised using unsupervised competitive learning driven by the standard trace learning rule (6). In their model the trace learning rule was able to exploit the natural statistics of how primates tend to move their eyes and head as they explore their visual environment, with more frequent shifts in eye-position than head position. These natural eye and head movements enable trace learning to bind together different visual input patterns corresponding to visual targets located in the same head-centred location but different retinal locations, thereby driving the development of postsynaptic neurons that respond to visual targets in specific head-centred locations.

It was originally demonstrated by [1], and shown again above, that the self-organising model described in section Materials and methods using the standard trace learning rule (6) with peaked eye-position gain modulated input neurons is able to develop head-centred output representations during training. However, a key new result in this paper is that the self-organising model with purely sigmoidal eye-position gain modulated input neurons develops eye-centred output neurons when using the standard trace learning rule. The cause of this contrasting behaviour between peaked and sigmoidal gain fields is the correlations between the activity of the input neurons across all of the input patterns during training. It is well understood that in a standard competitive neural network, individual output neurons learn to respond to subsets of input neurons that tend to be most frequently co-active [18]. In the self-organising model with peaked gain modulation, the subsets of input neurons that are frequently co-active correspond to circular clusters that are highly localised in both the retinotopic preference and eye-position preference dimensions. With a standard hebbian learning rule with no significant memory trace of recent neuronal activity, individual output neurons will learn to respond to these localised circular clusters of input neurons. If a trace learning rule is implemented, it is a relatively easy task for individual output neurons to simply bind together these clusters of input neurons along a diagonal line in the input space corresponding to a particular head-centred location. Output neurons will then respond to particular head-centred locations regardless of eye-position or the retinal location of a visual target. However, the situation is quite different with sigmoidal gain modulated input neurons. Due to monotonic saturation of the input neuron response function in the eye-position dimension of the input space, the subsets of input neurons that are most frequently co-active are localised in the retinotopic preference dimension but elongated in the eye-position preference dimension. With a standard hebbian learning rule, individual output neurons will learn to respond to these elongated clusters of input neurons, which will give rise to eye-centred output responses. However, if a trace learning rule is implemented, the output neurons still learn to represent eye-centred rather than head-centred locations. This is because developing head-centred output responses would require the trace learning rule to disrupt and break apart output representations corresponding to the elongated clusters of input neurons with correlated activities. However, in practice the trace learning effect is not strong enough to achieve this. Consequently, even with trace learning, the elongated clusters of input neurons with correlated activities continue to drive the development of eye-centred output neurons. This finding holds for any input neuron response function with a monotonic gain in the eye-position dimension, be it purely linear [14] or linear rectified [20].

We next showed that a manually prewired neural network model with sigmoidal eye-position gain modulated input neurons could perform the coordinate transformation. This is achieved by elevating the weight of synaptic connections from all input neurons which responded strongly, for some eye-position, to a visual target in the head-centred location to which the postsynaptic output neuron is assigned. This result established the feasability of the coordinate transformation within the given model architecture with sigmoidal gain fields acting on the input population. However, when synaptic plasticity based on the standard trace learning rule (6) is suddenly introduced into the manually prewired model, it is found that just a single epoch of visually-guided training is sufficient to dramatically reduce the prevalence of head-centred neurons in the output population, and by the 5^th^ epoch all neurons are eye-centred for all subsequent epochs of training. This showed that sigmoidal gain modulation would, in so far as the synapses are plastic, even undermine the functioning of a cortical circuit which is somehow prewired, perhaps by genetic hardwiring or by an initial period of supervised learning as suggested by [20], to produce the head-centred visual representations. Given the ubiquitous presence of associative synaptic plasticity in cortex, it is therefore a substantial challenge to explain how head-centred visual neurons might develop and persist in the brain given the relatively large proportion of precursor eye-centred visual neurons that have monotonic eye-position gain fields.

It has been shown that neurons in many parietal areas have a mixture of different forms of gain modulation, not all of which are planar. In fact, 41% and 56% are not planar in areas LIP and 7a, respectively [8]. Consequently, we explored how various degrees of prevalence of sigmoidal gain modulation would influence the self-organisation of the model with the standard trace learning rule (6). It is found that so long as there is no more than approximately 20% sigmoidal gain modulation in the input population then the fraction of head-centred neurons in the trained model did not drop below ~15%. However, the decrease in performance, and eventual collapse of the model, is very severe when the presence of sigmoidal gain modulation in the input population is increased. In particular, with a more biologically realistic 50:50 mix in peaked and sigmoidal gain modulated input neurons, the model failed to develop a significant population of head-centred output neurons with the standard trace learning rule (6).

In order to find a biologically plausible way in which the self-organising model could develop relatively large numbers of head-centred visual output neurons when at least half or even all of the input neurons had sigmoidal gain fields, we next explored the performance of the model architecture using a variety of more sophisticated learning rules that may incorporate temporally delayed memory traces, as well as a mixture of both trace learning and anti-Hebbian learning terms. These new, modified learning rules were previously developed by [25] in the context of modelling transform invariant visual object recognition. The new learning rules continue to be biologically plausible because they rely on locally available biological quantities, namely the activities of the pre- and post-synaptic neurons, to guide the modification of synaptic weights. The different modified learning rules were found to successfully drive the self-organisation of head-centred output responses when all of the input neurons had sigmoidal eye-position modulation. The modified learning rules were able to substantially improve the temporal binding of the standard trace learning rule by incorporating a trace of previous neuronal activity with an explicit time delay Δ*T*. This was evidenced by better performance being achieved when the learning rule incorporated a trace value of the postsynaptic neuron from an earlier time Δ*T* in the past instead of the trace value at the current time. Furthermore, the new learning rules were able to overcome the high correlations between overlapping input patterns with sigmoidal gain fields through the additional incorporation of anti-Hebbian learning, which had been previously found to offer a significant performance enhancement by [25]. However, learning rule (20) incorporated a delayed postsynaptic trace without an anti-Hebbian term (section Modified Learning Rule: Delayed Postsynaptic Trace Learning Rule). Simulation results with this learning rule showed that head-centred output responses developed in the trained model. Thus, anti-Hebbian learning is not necessarily a requirement for the development of head-centred output responses. The performance of learning rule (20) was then investigated under a more biologically realistic scenario in which the input population consisted of a 50:50 mixture of neurons with either peaked or sigmoidal eye-position gain modulation. These results demonstrated that the model could produce large numbers of head-centred output neurons with a learning rule that relied purely on a postsynaptic delayed-trace term without any anti-hebbian term if the input population contained a 50:50 mix of neurons modulated by either peaked or sigmoidal gain fields. These last findings should be contrasted with the performance of
the model with the standard trace learning rule (6), which failed to develop a significant 1251 number of head-centred output neurons with such a mixed input population.

## Conclusion

In conclusion, the work presented here has shown that the existence of a synaptic weight structure that can perform a coordinate transformation does not guarantee that it will emerge through a process of self-organisation using any one particular form of trace learning rule. In particular, the standard trace learning rule originally proposed by [1] failed to produce head-centred output neurons when the input neurons were modulated by a sigmoidal function of eye-position, and even failed with a biologically realistic 50:50 mix of input neurons with peaked and sigmoidal gain fields. In order to remedy this problem, we had to investigate the use of more sophisticated, yet still biologically plausible, learning rules that incorporated temporally delayed memory traces, as well as a mixture of both trace learning and anti-hebbian learning terms. These new, modified learning rules were found to produce head-centred output neurons when the input population had sigmoidal gain fields. Moreover, it was also found that the delayed postsynaptic trace in learning rule (20) was sufficient to drive the development of head-centred output neurons in the absence of anti-hebbian learning, especially if the input population had a 50:50 mix of peaked and monotonic gain fields. This work thus makes an important contribution to understanding how head-centred visual neurons may develop in the brain through an unsupervised process of visually-guided learning given visual precursor neurons with sigmoidal (monotonic) eye-position modulation. Furthermore, although we have studied one particular kind of coordinate transformation, i.e. from eye-centred to head-centred visual representations, these findings may also apply to a range of other coordinate transformations with sensorimotor integration of monotonically encoded motor variables [4,5].

## Supporting information

**Appendix A. Eye-centredness Reference Frame Analysis.** During testing, the visual target was located in head-centred locations

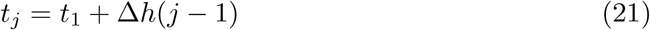

for *j* = 1, …, *T*, and while in each location it was observed from eye-positions

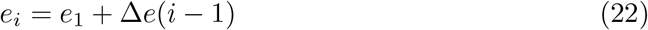

for *i* = 1, …, *E*. The eye-position shift Δ*e* was set to a multiple of the head-centred target location shift Δ*h* to cause resampling of the neuron’s response at the same retinal location for different eye-positions, thereby providing a resampling of the response in both head-centred and eye-centred space across multiple eye-positions.

The set of head-centred locations {*t*_1_,…,*t_T_*} corresponded to retinal locations *R_i_* = {*t*_1_ − *e_i_*,…,*t_T_* − *e_i_*} when the model was fixating eye-position *e_i_*, and from this it is clear that among retinal locations common to all eye-positions, *t_1_* − *e*_1_ was the first and *t_T_* − *e_E_* was the last, that is

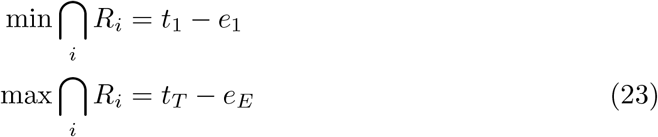

Therefore *f_i_* and *l_i_,* denoting the first and last position included from the *i*^th^ response vector respectively, had to correspond to these two retinal locations respectively

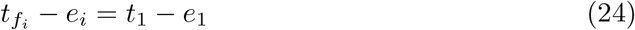

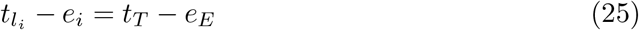

We can resolve each equation to find an explicit formula for *f*_*i*_ and *l*_*i*_ in terms of *i* as follows.

By substituting Eq 21 and Eq 22 into Eq 24 we obtain

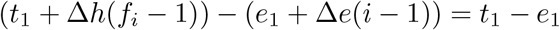

Rearranging this gives

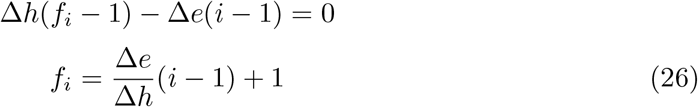

By substituting Eq 21 and Eq 22 into Eq 25 we obtain

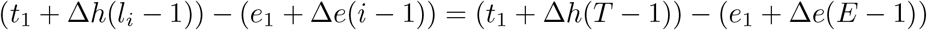

Rearranging this gives

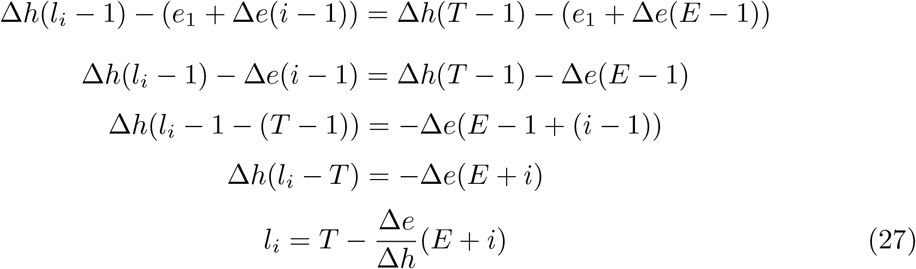

We can also deduce the length *V* of the portion of each response vector that is used in the eye-centred correlation analysis as follows. By definition, for each response vector

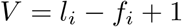

Substituting in Eq 26 and Eq 27 gives

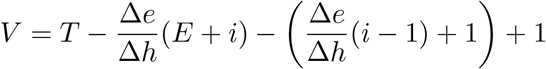

Rearranging gives

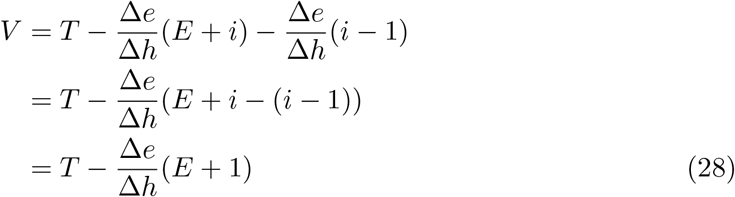

## Acknowledgments

This work was supported by CNPq, National Council for Scientific and Technological Development - Brazil.

